# Predictable clonal hierarchies from restricted progenitors provide a framework for cell type-specific therapies in glioblastoma

**DOI:** 10.64898/2026.02.21.707071

**Authors:** Elisa Fazzari, Daria J. Azizad, Matthew X. Li, Weihong Ge, Shivani Baisiwala, Dimitri Cadet, Patricia R. Nano, Ryan L. Kan, Travis Perryman, Hong A. Tum, Christopher Tse, Brittney Wick, Carolina Varona Arguelles, Kunal S. Patel, Linda M. Liau, Robert M. Prins, David A. Nathanson, Aparna Bhaduri

## Abstract

Extensive molecular profiling has revealed profound heterogeneity in glioblastoma (GBM), yet how cellular lineages organize over time to govern tumor propagation and therapeutic response remains poorly understood. Existing single-cell approaches define transcriptional states but provide limited insight into how clonal dynamics shape functional tumor behavior. Here, we integrate high-complexity combinatorial DNA barcoding with single-cell transcriptomics in direct-from-patient IDH1-wild-type GBM, enabling lineage-resolved mapping of progenitor organization in a human microenvironmental context. Across 235,155 malignant cells from nine tumors, clonal relationships form reproducible lineage tracks in which distinct progenitor populations give rise to specific differentiated cell types, revealing that tumor growth is sustained by multiple non-redundant progenitors rather than a single dominant population. These progenitors exhibit distinct propensities for self-renewal, fate restriction, and cross-compartment interactions, collectively accounting for the full spectrum of tumor states. Using this lineage-resolved framework, we identify complementary drug targets in distinct progenitor compartments and demonstrate that hierarchy-informed combination therapies disrupt progenitor-progenitor interactions and reshape lineage output. These findings move beyond descriptive heterogeneity to define functional logic underlying GBM propagation and establish a generalizable framework for rational, cell type-specific combinatorial therapies.

## Introduction

Glioblastoma (GBM) exhibits profound cellular heterogeneity, yet how this diversity is functionally organized into progenitor hierarchies and exploited therapeutically remains unclear. Two prevailing models have emerged: one posits that GBM growth is organized by invariant stem cell hierarchies^1^; the other proposes that malignant states exist in a largely plastic equilibrium, with frequent interconversion between transcriptionally defined cell states^2^. This hierarchical framework is further refined by cell-of-origin identity: identical tumor suppressor mutations in distinct adult CNS progenitor populations generate molecularly and biologically distinct GBM subtypes, each retaining transcriptional signatures of their lineage of origin and exhibiting unique therapeutic vulnerabilities^3,4^. At the same time, tumor cell states can also undergo dynamic transitions, including therapy-associated proneural-to-mesenchymal shifts and reversible persister programs^5^, while epigenetic regulatory architectures constrain or permit these transitions^6^. Lineage barcoding has further indicated that genetic and epigenetic mechanisms cooperate to drive therapeutic resistance^7^.

Single-cell and spatial profiling studies have converged on the view that GBM tumors contain multiple transcriptionally and functionally distinct populations that behave as progenitors rather than a single homogeneous stem cell compartment. Radial glia-like cells (RG)^8^, outer radial glia (oRG)^9^, oligodendrocyte progenitor-like cells (OPC), neural progenitor-like cells (NPC)^4,10^, and intermediate progenitor populations with tri-lineage potential (Tri-IPC)^11^ have all been independently identified across patient cohorts and analytical frameworks^2,8–10,12,13^. In this study, we use the term progenitor to describe these developmentally informed, transcriptionally defined tumor cell populations that express neural stem or precursor-associated gene programs and exhibit proliferative features distinct from terminally differentiated tumor states. Lineage tracing and single-cell studies further indicate that these populations differ in proliferative dynamics and responses to therapeutic pressure, including the ability to enter drug-tolerant persister states or undergo compensatory reprogramming^7,14^. Many of these progenitor populations resemble cell types active during human cortical development, underscoring the extent to which GBM reactivates neurodevelopmental programs. In particular, oRG-like cells, which are evolutionarily expanded in the human brain, have emerged as a recurrent population in GBM with enhanced migratory capacity and lineage plasticity^8,9,15^. The prominence of such human-enriched progenitor states highlights the importance of studying tumor hierarchies in human-specific systems and supports a growing paradigm in which malignant progression reflects the reactivation and recombination of developmental lineage programs^2,10^.

In parallel, large-scale single-cell atlases have repeatedly identified overlapping progenitor and differentiated states, providing confidence that these populations represent conserved features of human disease rather than study-specific artifacts^2,9,16–20^. These atlases now poise the field to functionally interrogate, in human tumor samples, how tumor heterogeneity drives tumor clonal expansion and how distinct progenitors generate all or some of the expanding tumor. Three fundamental questions remain unresolved. First, do transcriptionally defined progenitor states correspond to durable functional differences in lineage restriction and plasticity, or do they primarily reflect reversible expression programs? Second, does the degree of plasticity within a tumor meaningfully influence its aggressiveness or therapeutic resistance-that is, does lineage restriction itself carry clinical consequence? Third, do progenitor compartments with distinct fate biases exhibit selective therapeutic vulnerabilities that can be rationally targeted without provoking compensatory reconstitution of the hierarchy?

Here, we address these questions by integrating high-complexity lineage tracing with single-cell transcriptomics in direct-from-patient GBM samples. We show that GBM hierarchies are neither strictly linear nor dominated by a single apex progenitor capable of generating all cellular states. Instead, distinct progenitors occupy characteristic positions within the hierarchy, exhibit unique patterns of fate restriction and cross-lineage interaction, and leave reproducible footprints on tumor composition. These properties help explain why strategies aimed at eliminating a single progenitor compartment often fail to produce durable responses, as remaining progenitors can sustain or reconstitute the hierarchy through compensatory mechanisms. We therefore propose a lineage-informed therapeutic design framework that emphasizes combinatorial targeting of functionally complementary progenitor populations and provide both a dataset and a proof-of-concept demonstration of how such treatments can be rationally designed and evaluated.

## Results

### Lineage tracing in primary GBM cells

Multiple organoid models of GBM are emerging as tractable ways to preserve tumor cell type diversity in a human microenvironmental context^21–24^. The human organoid tumor transplant model (HOTT)^25^ encompasses these strengths while enabling use of direct-from-patient primary resected tumors and providing molecular access for lineage barcode delivery. We leveraged this system to explore the functional consequences of tumor heterogeneity in direct-from patient GBM samples (n=9). The CellTagging system^26^ enables a high degree of complexity by leveraging a combinatorial barcoding strategy in which a cell’s unique combination of barcodes defines clonal relationships. This innovation enables the study of direct-from-patient cells without prior *in vitro* expansion, preserving nuanced tumor-progenitor functions that are lost in conventional *in vitro* systems. CellTagged cells are visualized in the HOTT model via GFP expression and are subsequently harvested for lineage reconstruction and cell identity profiling via single-cell RNA-sequencing (scRNA-seq).

Tumor samples were restricted to untreated, newly diagnosed GBM (IDH1^WT^) and encompassed a variety of expected genetic profiles (SFig. 1a). Patient tumor samples were dissociated into single-cell suspensions and transduced with the CellTag lentiviral barcoding library at a target MOI of 3-4 barcodes per cell. In order to avoid *in vitro* selection processes, cells were infected for only 60 minutes and directly transplanted onto human cortical organoids (Fig. 1a, Methods). During a proliferation period of about two weeks, tumor cells were monitored to visualize invasion and morphological complexity (Fig. 1a-b, SFig. 1b). At the endpoint, transplants were dissociated and GFP+ tumor cells were harvested by fluorescence-activated cell sorting (FACS). GFP+ tumor cells were then captured for scRNA-seq (Fig. 1a-b, SFig. 1b-c, Methods). scRNA-seq data was subject to stringent quality control (Methods). Although only tumor cells received CellTag viral infection and thus GFP labeling, we further filtered our data to ensure only malignant cells were included based upon copy number variation analysis (CNV) due to recent reports that fluorescent protein transfer can occur between tumor cells and the microenvironment but that CNVs are restricted to tumor cells^22^.

**Figure 1.**
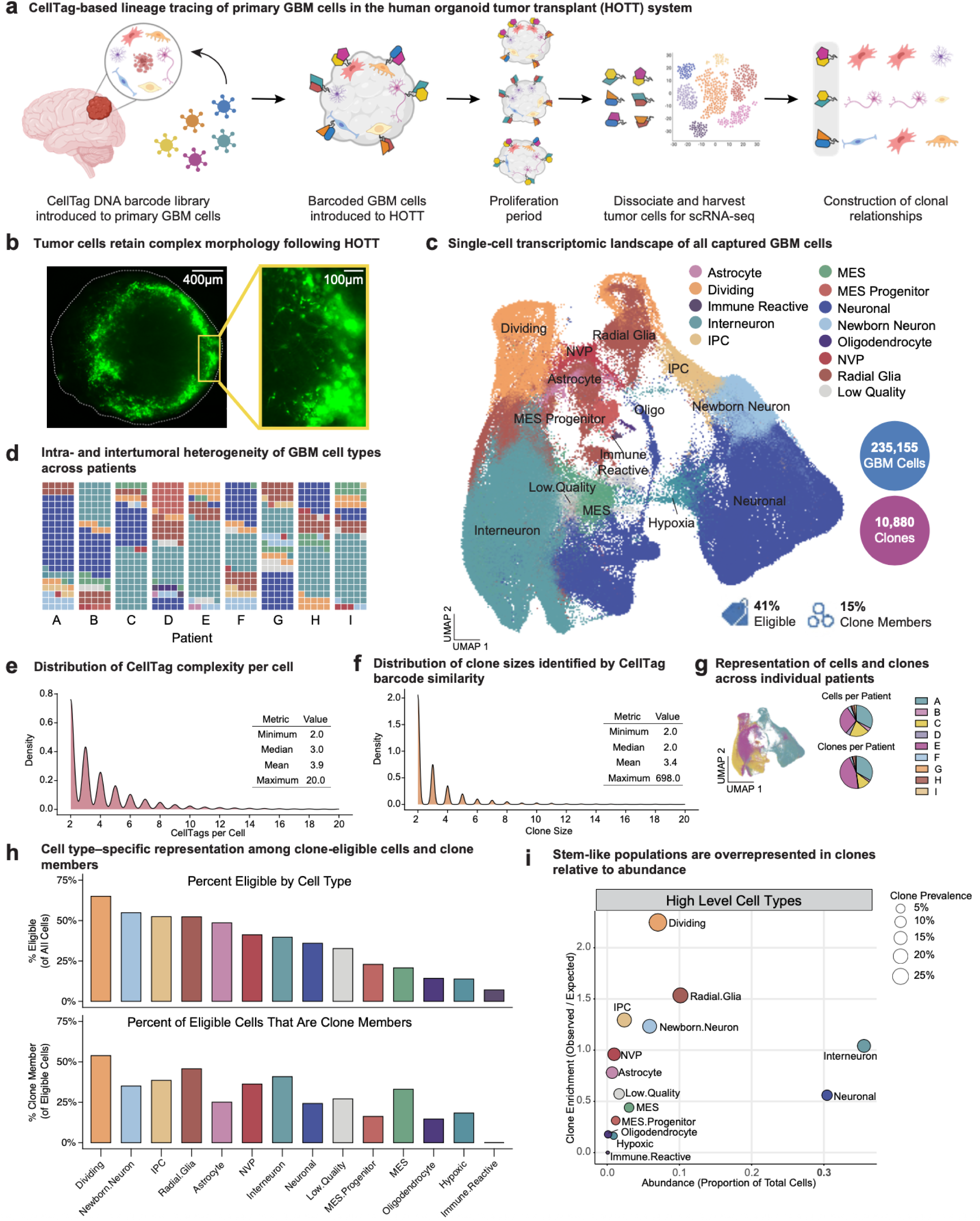
CellTag-based lineage tracing in a human organoid tumor transplant system enables clonal analysis of primary glioblastoma a) Schematic of the CellTag-based lineage tracing workflow in the human organoid tumor transplant (HOTT) system. Primary Isocitrate Dehydrogenase 1 wild-type (IDH1^WT^) glioblastoma (GBM) cells are dissociated and transduced with a high-complexity CellTag lentiviral barcode library encoding combinatorial DNA barcodes and GFP. Barcoded tumor cells are directly transplanted onto week 8-12 human cortical organoids^25^, allowed to proliferate without prior *in vitro* expansion, then dissociated for single-cell RNA sequencing (scRNA-seq). Lineage relationships are reconstructed by identifying shared CellTag barcode combinations across cells. b) Representative fluorescence images showing GFP⁺ live GBM cells within cortical organoids following transplantation, illustrating complex tumor morphology and invasion into the organoid tissue. Left, low-magnification overview of a transplanted organoid; right, higher-magnification inset highlighting heterogeneous tumor cell morphologies. Scale bars, 400 µm (left) and 100 µm (right). c) Uniform Manifold Approximation and Projection (UMAP) visualization of 235,155 quality-controlled malignant GBM cells captured across (n = 9) primary IDH1^WT^ patient tumors following CellTag-based lineage tracing in the HOTT system. Cells are colored by transcriptional cell type or state annotation derived using a multi-pronged strategy integrating (i) label transfer from a GBM meta-atlas of primary tumors using Seurat MapQuery^49^, (ii) scoring of published GBM cell state gene modules^2,18^, and (iii) cluster marker analysis informed by human neurodevelopmental reference datasets^53,54^. Annotated malignant cell populations include progenitor populations: Radial Glia (RG), Mesenchymal Progenitors (MES.Progenitor), Intermediate Progenitor Cells (IPC), and an Astrocyte population with progenitor-like features (Astrocyte), and Neurovascular Progenitors (NVP), as well as differentiated or state-defined tumor populations including Neuronal (Neuronal), Newborn Neurons, Interneuron, Oligodendrocyte-like (Oligo), Mesenchymal (MES), Immune Reactive, Hypoxic (Hypoxia), and Dividing cells. Of the total malignant cells, 41% met stringent CellTag complexity and quality criteria and were eligible for clone calling. Among all captured cells, 15% were assigned as clone members, yielding a total of 10,880 inferred clones across the dataset. d) Intra- and intertumoral heterogeneity of GBM cell type composition across individual patients. Waffle plots display the relative abundance of annotated transcriptional cell types for each of the nine patient tumors profiled in the HOTT system. Each square represents 1% of the total malignant cell population for a given patient and is colored by cell type as defined in panel c. This representation highlights both shared and patient-specific differences in tumor cell state composition across patients. e) Distribution of CellTag barcode complexity per cell among clone-eligible tumor cells. Density plot shows the number of CellTags detected per cell after filtering to retain cells with sufficient barcode complexity for downstream clone calling (2-20 CellTags). Summary statistics (minimum, median, mean, maximum) are calculated across all clone-eligible cells and displayed as an inset table. f) Distribution of inferred clone sizes across tumors. Density plot depicts clone size (number of cells per clone) for clones identified by CellTag barcode similarity, plotted from the ground-truth clone size table and visualized over the range shown (2-20 cells on the x-axis). Summary statistics (minimum, median, mean, maximum) are calculated from the full clone-size distribution and shown as an inset table. g) Representation of malignant GBM cells and inferred clones across individual patients. Left, UMAP projection of all quality-controlled malignant cells colored by patient of origin, illustrating intermixing of patient-derived cells in transcriptional space. Right, pie charts showing the proportion of total cells (top) and inferred clones (bottom) contributed by each patient. h) Cell type-specific representation among clone-eligible cells and clone members. Top, percentage of all cells within each annotated cell type that meet stringent criteria for clone calling, calculated with the total number of cells of that type as the denominator. Bottom, percentage of clone-eligible cells within each cell type that are assigned as members of inferred clones, calculated using clone-eligible cells of that type as the denominator. Cell types are colored consistently with panel c. i) Clone enrichment versus abundance across high-level GBM cell types. Bubble plot summarizes, for each annotated cell type, its overall abundance in the dataset (x-axis; proportion of all malignant cells) and its relative enrichment among clone members (y-axis; observed/expected). Clone enrichment is calculated as the fraction of all clone-member cells assigned to a given cell type divided by that cell type’s fraction of all cells: (Clone Member Count/Total Clone Members)/(CellType Count/Total Cells). Bubble size reflects clone prevalence, defined as the fraction of cells within a given cell type that are clone members (Clone Member Count/CellType Count). Points are colored by cell type. Values >1 indicate overrepresentation among clone members relative to abundance, whereas values <1 indicate depletion.

After quality control measures, the resulting dataset consisted of 235,155 tumor cells across the 9 patient tumor specimens (Fig. 1c-d, 1f). In a companion study^27^, we leveraged a meta-analysis of published scRNA-seq data to annotate the heterogeneity of cell types that exist in GBM, informed by neurodevelopmental analogs. We accordingly annotated our cell types from our CellTagged cells with these states, and in some cases combined subcategories of similar cell types (Fig. 1c-d, SFig. 1d, STable2). Putative progenitor populations included radial glia^8,9^, mesenchymal progenitors (MES.Progenitor), intermediate progenitor cells (IPC)^11^, a dual-fate progenitor called the Neurovascular Progenitor (NVP)^27^ and an astrocyte population exhibiting progenitor-like features^2,18,28,29^, consistent with prior observations of proliferative and lineage-competent astrocytic states in GBM. In addition to these putative progenitors, we identified presumed differentiated or populations including neurons, newborn neurons, interneurons, oligodendrocytes (Oligo), immune-reactive^30^ tumor cells, mesenchymal (MES) cells, and hypoxic^18,31^ tumor cells. Across the 9 patients, we observed intra- and intertumoral heterogeneity as previously described^2,9,18,19,30–32^ with most cell types represented across most patients (Fig. 1d, 1g).

Independently and orthogonally to cell type annotation, the quality-controlled tumor cells were then subject to CellTag barcode analysis in order to identify clones within our dataset. Clone analysis was performed within each tumor sample individually. We observed 41% of cells met the criteria to be included in clone calling (Fig. 1c,e-f, SFig. 1f-g, STable3, Methods), with a set range of 2 to 20 barcodes per cell and average of 3.9. Using a simulation analysis, we verified that the probability of two cells being determined as clonally related due to spurious labeling was infinitesimally small (Methods). Determining clone membership was based upon the jaccard correlation of CellTags, and the clone analysis resulted in 10,880 clones with an average clone size of 3.4 cells (Fig. 1c,e,f,g, SFig. 1f-g, Methods).

After cell type annotation and clone calling were individually completed (SFig. 1f, STable4, Methods), clone information was mapped onto cell types to functionally interpret the data. Both the fraction of cells eligible for clone calling and clone membership were enriched across populations that have been described as having stem-like properties in GBM^8,9,11,29^. We proceeded to explore the relationship between cell type abundance and likelihood of that cell type to be represented in clones (Fig. 1h-i, SFig. 1g). As expected, dividing cells and radial glia populations were overrepresented in clones compared to their abundance. Key negative controls such as the immune reactive and hypoxic populations had little to no representation in clones. Similarly, we observed a high abundance of neuronal and interneuron populations which is consistent with prior descriptions of the HOTT system^25^, but these were not overenriched in clones. We observed that our annotated states offered more granularity beyond widely used GBM cell states and aligned well to these published studies^2,18^ (SFig. 1d-e).

### Clonal relationships among glioblastoma cell types organize into track-like groupings

Upon evaluation of individual clones determined by barcode similarity, we noted that 34% of clones contained only one cell type (SFig. 2a-b), suggesting a large fraction of symmetrical divisions. To decipher clonal relationships that drive tumor propagation, we further explored the patterns that emerged from multi-cell type clones including those that recapitulate previous GBM and neurodevelopmental literature (Fig. 2a). For example, we observed canonical excitatory (Fig. 2ai) and putative inhibitory (ii) neurodevelopmental trajectories consistent with previously described reactivation of neurodevelopmental programs in GBM. Specifically, we found that outer radial glia (oRG) were clonally related to diverse GBM neuronal subtypes^8–10^ (iii). We also noted clonal relationships between oRGs, excitatory neurons and interneurons (iv) which is consistent with recent lineage tracing studies of interneuron generation in the developing human cortex^33^. Additionally, we confirmed the existence of a predicted tripotent IPC (tri-IPC) identified in human cortical development and hypothesized to exist in GBM (v-vi)^11^. Strikingly, we identified a subpopulation with relationships in both the neural and mesenchymal lineages (vii-viii) that we named a neurovascular progenitor (NVP). A companion study showed that NVP cells exist in primary GBM and exhibit transcriptional profiles of both vascular, perivascular-like cells and neural progenitor cells^27^.

**Figure 2:**
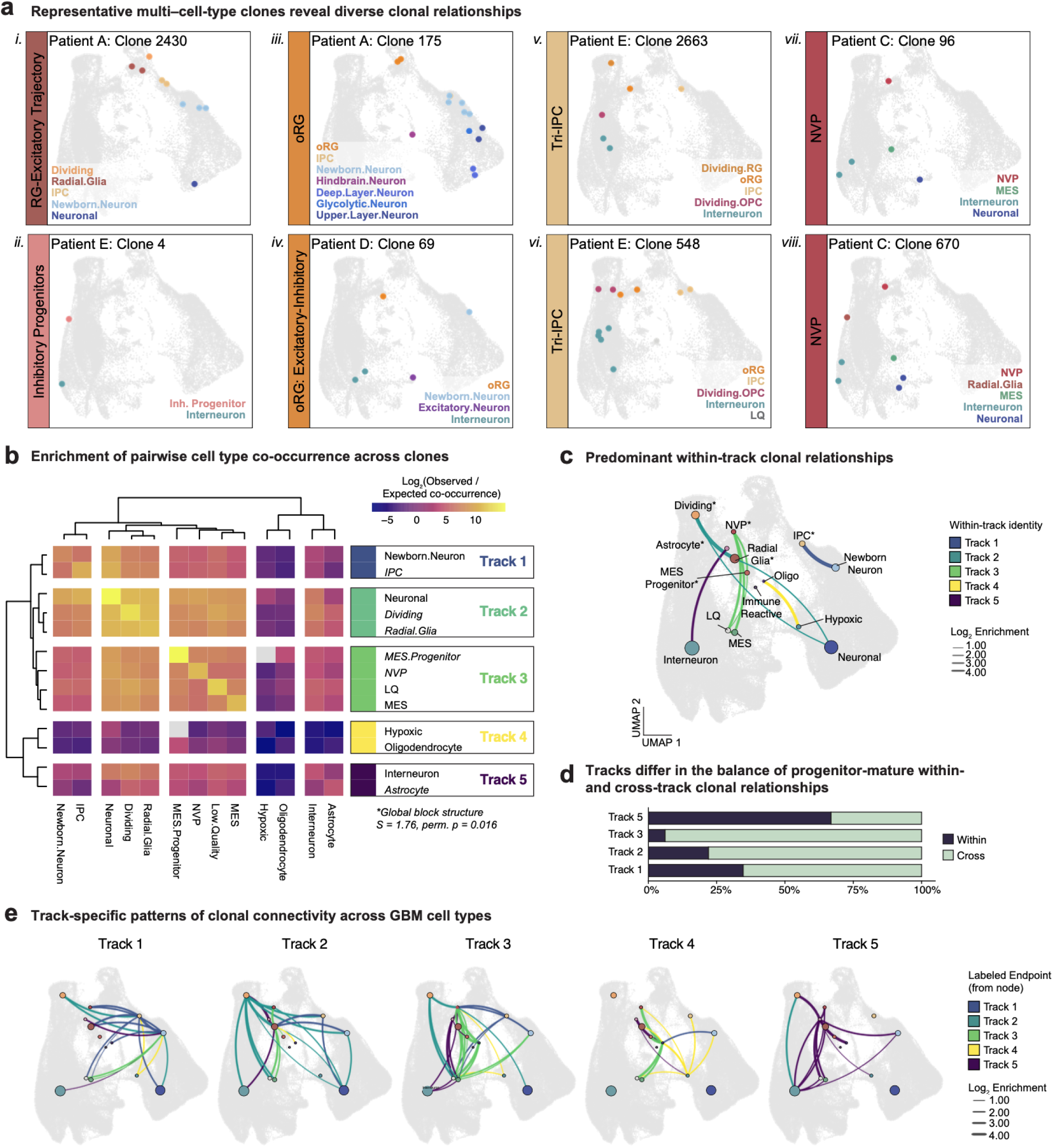
Clonal relationships among GBM cell types organize into track-like groupings a) Representative multi-cell-type clones illustrating diverse clonal relationships among GBM cell types. Each panel shows a single inferred clone overlaid on the global UMAP (gray), with colored points indicating clone member cells colored by cell type. Individual clones highlight canonical radial glia-associated excitatory trajectories (i), putative inhibitory progenitor trajectories (ii), outer radial glia (oRG) clones with clonal links to multiple neuronal subtypes (iii), oRG clones exhibiting both excitatory and inhibitory fates (iv), predicted tripotent intermediate progenitor cell (Tri-IPC) clones (v-vi), and neurovascular progenitor (NVP) clones exhibiting relationships spanning neural and mesenchymal lineages (vii-viii). b) Enrichment of pairwise cell type co-occurrence across clones. Heatmap shows the log₂(observed/expected) enrichment of cell type pairs co-occurring within the same inferred clone. Progenitor populations are indicated in italics. Observed co-occurrence was calculated as the number of clones containing both cell types, while expected co-occurrence was derived from a null model based on the overall abundance of each cell type across all malignant cells and the total number of clones (Methods; Supplementary Fig. 2e). Positive values indicate cell type pairs that co-occur in clones more frequently than expected by chance, whereas negative values indicate underrepresentation. Hierarchical clustering of the enrichment matrix reveals five major communities of cell types, termed Tracks (Tracks 1-5), representing groups of cell types that preferentially co-occur across clones. Block structure significance was quantified using a global blockiness statistic, defined as the difference between the mean absolute enrichment within dendrogram-defined blocks and between blocks, and assessed by permutation of cell type labels (S = 1.76, permutation p = 0.016), indicating non-random organization of clonal relationships. c) Network visualization of clone-level co-occurrence among GBM cell types restricted to within-track interactions. Each node represents a cell type positioned at the median UMAP coordinates of its constituent cells, with node size proportional to the total number of cells assigned to that type. Edges denote significant enrichment of co-occurrence within clones relative to expectation based on marginal cell type frequencies, with edge thickness scaled by the log₂(observed/expected) enrichment score. Edge color indicates the developmental track identity of the originating cell type, as defined by track assignments in Fig. 2b. d) Stacked bar plots show, for each clonal Track, the relative fraction of enriched progenitor-mature cell type relationships occurring within the same Track compared with those spanning different Tracks. e) Each panel displays enriched clone-level co-occurrence relationships involving cell types assigned to the indicated developmental Track. Edges denote statistically enriched clonal co-occurrence between cell type pairs (log₂ observed/expected ≥ 0.5), indicating that the two cell types co-occur within the same clone more frequently than expected based on marginal frequencies. Edge thickness reflects the magnitude of log₂ enrichment. For each panel, edges are shown if at least one endpoint belongs to the indicated Track. Edge color corresponds to the developmental Track assignment of the labeled endpoint (from node).

We next asked what patterns emerge across all of the clones observed in our dataset while being mindful that cell types exist in different proportions in patient tumors. We therefore calculated the expected co-occurrence of cell type pairs in clones given clone size and cell type abundance and investigated the observed enrichment of pairwise clone relationships across our dataset (SFig. 2e, Methods). We found that clone relationships were divided into five major hierarchies, which we call Tracks.

Inspection of the global co-occurrence structure revealed that the five Tracks correspond to biologically coherent groupings of cell types, each organized around a progenitor-like population (*italicized* populations in Fig 2b.) that anchored clonal relationships within that Track. Track 2 captured a continuum spanning radial glia, dividing, and neuronal populations, consistent with a canonical radial glia-to-neuronal lineage relationship widely observed in both developmental neurobiology and proneural GBM programs (Fig. 2b-e). In contrast, Track 3 encompassed MES.progenitor, NVP, and MES populations, aligning with a mesenchymal hierarchy. Notably, Track 3 displayed extensive cross-Track relationships, indicating that putative mesenchymal-associated progenitors participate broadly in clonal interactions beyond a single lineage compartment. Track 1 represented a grouping composed of intermediate progenitor cells (IPC) and newborn neurons. The tight association between these populations suggests a transitional clonal module linking presumed progenitor and early neuronal identities, consistent with developmental IPC-newborn neuron relationships Track 5, consisting of interneurons and astrocytes, was distinguished by strong within-Track co-occurrence and minimal connectivity to other Tracks, indicating comparatively rigid clonal constraints. Although astrocytes are often considered differentiated end states, accumulating evidence supports progenitor-like and proliferative properties in subsets of tumor-associated astrocytes^28,29^, consistent with their placement as a Track-defining population here (Fig. 2b, SFig. 2c). Finally, Track 4, which contained hypoxic cells and oligodendrocytes, showed limited internal and external connectivity, suggesting that these states may reflect environmental or differentiation endpoints with reduced clonal mixing.

We hypothesized that tracks represent communities of cell types frequently found together across clones, and that they likely reflect broad axes of clonal proximity rather than strict lineages (Fig. 2b-e). In other words, Tracks are the data-driven groupings of cell type relationships, while different cell types have differing propensities to remain within or move between Tracks. After defining the major tracks, we interrogated which of them represent more rigid clonal constraints and which exhibit the most flexibility. Track 5, composed of interneurons and astrocytes, exhibits the majority of its relationships within cell types of the same track, while Track 3 included NVP and MES.Progenitors and exhibited nearly all of its relationships between cell types of different tracks (Fig. 2d-e). Although the relative strength and connectivity of individual Tracks varied across patients, with all tumors containing representatives of each Track, indicating a conserved clonal organization despite inter-patient heterogeneity (SFig. 3a). Stratification by EGFR expression further revealed modest shifts in Track connectivity between EGFR-high and EGFR-low tumors, suggesting that tumor genetics may subtly bias clonal relationships without altering the overall Track architecture (SFig. 3b).

**Figure 3.**
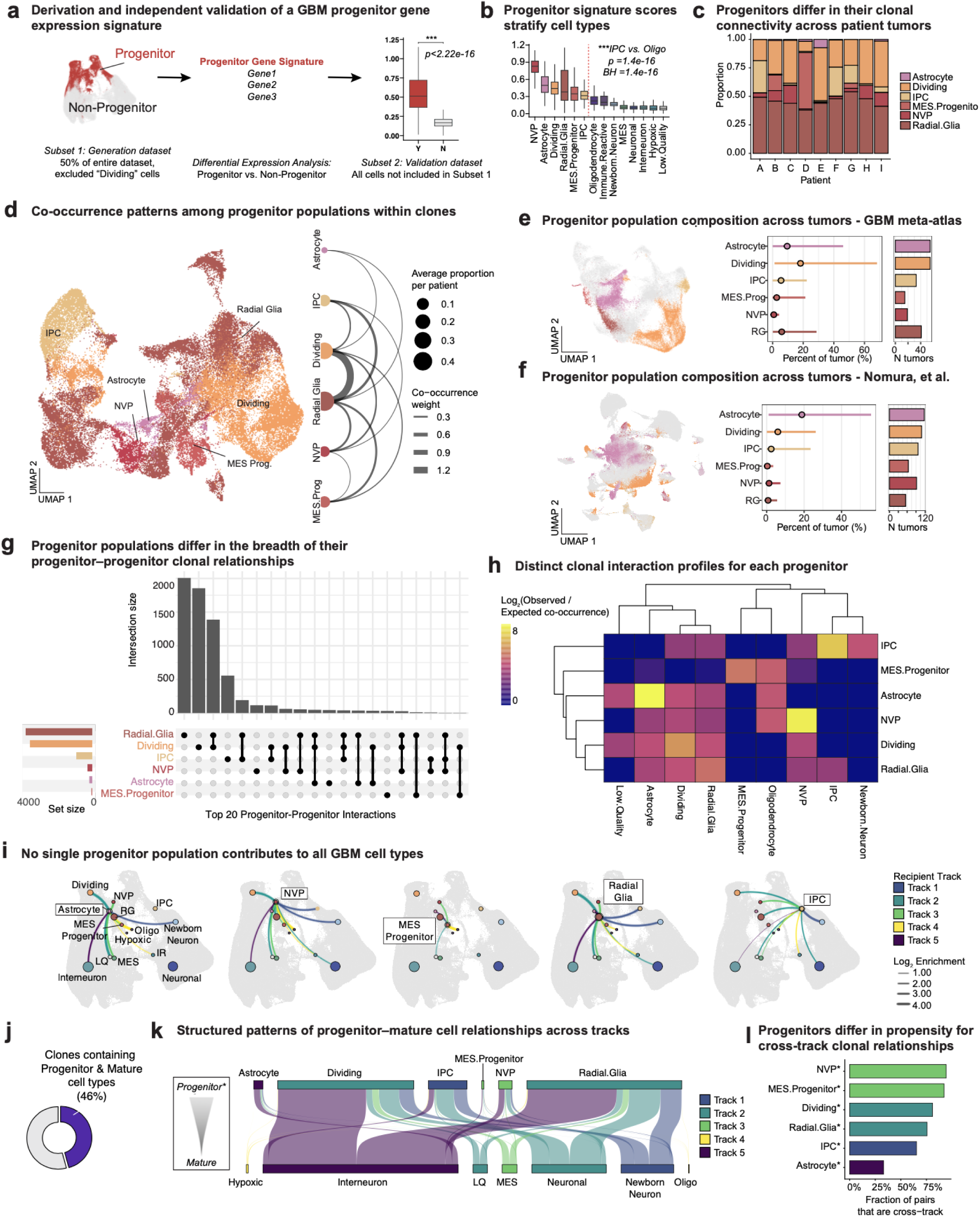
Progenitor-progenitor and progenitor-mature clonal relationships organize GBM cell type transitions a) Schematic of progenitor gene signature derivation and independent validation. The dataset was randomly split in half; a progenitor module was defined in the discovery subset (50% of cells; dividing cells excluded) by differential expression between progenitor-annotated cells and all other malignant cells, and then scored in the held-out validation subset (remaining 50%) using Seurat AddModuleScore. Boxplots show progenitor module scores in validation cells labeled as progenitor (Y) versus non-progenitor (N); significance was assessed by two-sided Wilcoxon rank-sum test (p value shown on plot). b) Progenitor module scores across transcriptional cell types, ordered by median score. Boxplots show the distribution of module scores per cell type in the validation set; the dashed red line indicates the progenitor-mature separation used in downstream analyses. Statistical comparisons were performed using pairwise two-sided Wilcoxon rank-sum tests with Benjamini-Hochberg correction for multiple testing; the example comparison shown is IPC versus Oligodendrocyte (BH-adjusted p value shown on plot). c) The composition of clone-member progenitor cells by progenitor identity (Radial.Glia, IPC, NVP, MES.Progenitor, Dividing, Astrocyte), expressed as proportions within each patient. d) Progenitor-only UMAP (left) and patient-weighted progenitor co-occurrence (right). Left, UMAP of the progenitor subset, colored by progenitor subtype (IPC, Radial Glia, Dividing, MES.Progenitor, NVP, Astrocyte). Right, arc diagram summarizing progenitor-progenitor co-occurrence across patients: node size denotes the average within-patient proportion of each progenitor subtype, and edge width reflects abundance-weighted co-occurrence (summed across patients as prop_A × prop_B for progenitor pairs exceeding a minimum within-patient proportion threshold; see Methods). e) Progenitor population composition across tumors in a GBM meta-atlas. Left, UMAP embedding with progenitor populations highlighted. Middle, distribution of the fraction of tumor cells comprised by each progenitor population across individual tumors. Right, proportion of tumors in which each progenitor population is detected. f) Same analyses as in e, performed on an independent GBM dataset from Nomura et al., with cell type labels assigned by MapQuery projection onto the GBM meta-atlas (Methods) g) Top 20 most frequent combinations of progenitor cell types co-occurring within individual clones containing at least one progenitor cell. Each column represents a unique combination of progenitor types (black dots), with bar height indicating the number of clones exhibiting that combination (intersection size). Horizontal bars indicate the total number of clones in which each progenitor type participates (set size). h) Progenitor-target co-clone enrichment patterns, with values computed using the same observed/expected framework described in the Methods and Supplementary Fig. 2e; for each progenitor, the top five positively enriched interactions are shown. i) All unique progenitor-mature cell pairs were enumerated within clones and classified as within-lineage or cross-lineage based on developmental track assignments. Network diagrams show enriched progenitor-mature relationships for each progenitor type, with edge thickness proportional to log₂ enrichment and color indicating recipient lineage. These patterns reveal biased but incomplete lineage outputs, demonstrating that no individual progenitor population gives rise to all GBM cell types. j) Fraction of clones containing both progenitor and mature cell types. k) Sankey diagram depicting clone-level relationships between progenitor cell types (top) and mature cell types (bottom). Flow width is proportional to co-occurrence across clones. Flows and target nodes are colored by the Track of the mature (target) cell type, indicating whether progenitor-mature relationships terminate within the same Track or span different Tracks. l) For each progenitor cell type, the fraction of all observed progenitor-mature cell pairs that span different Tracks (cross-track), calculated at the pair level across all clones.

### Progenitor identity shapes clonal fate, restriction, and cross-lineage interactions in GBM

A notable observation was that data-driven groupings of clone relationships tended to include a putative progenitor cell as defined by parallel function in the context of neurodevelopment (e.g. IPC, Radial Glia, etc) as well as a cell type traditionally thought of as more mature (e.g. Interneuron, MES). In order to more closely interrogate the hypothesis that these are in fact progenitor types, we derived a progenitor gene expression signature by subsetting our dataset to half and performing differential expression analysis between the putative progenitor cells and all other cells (Fig. 3a, STable5, Methods). We confirmed with gene ontology enrichment analysis that this signature mapped well to described progenitor signatures in the literature (Supp. Fig 4a). The other half of the dataset was used as a validation of our signature, in which all cells were scored for the gene expression program. Using our validation set, we identified which cell types had significant enrichment for our expression module compared to all other cells (Fig. 3a).

**Figure 4.**
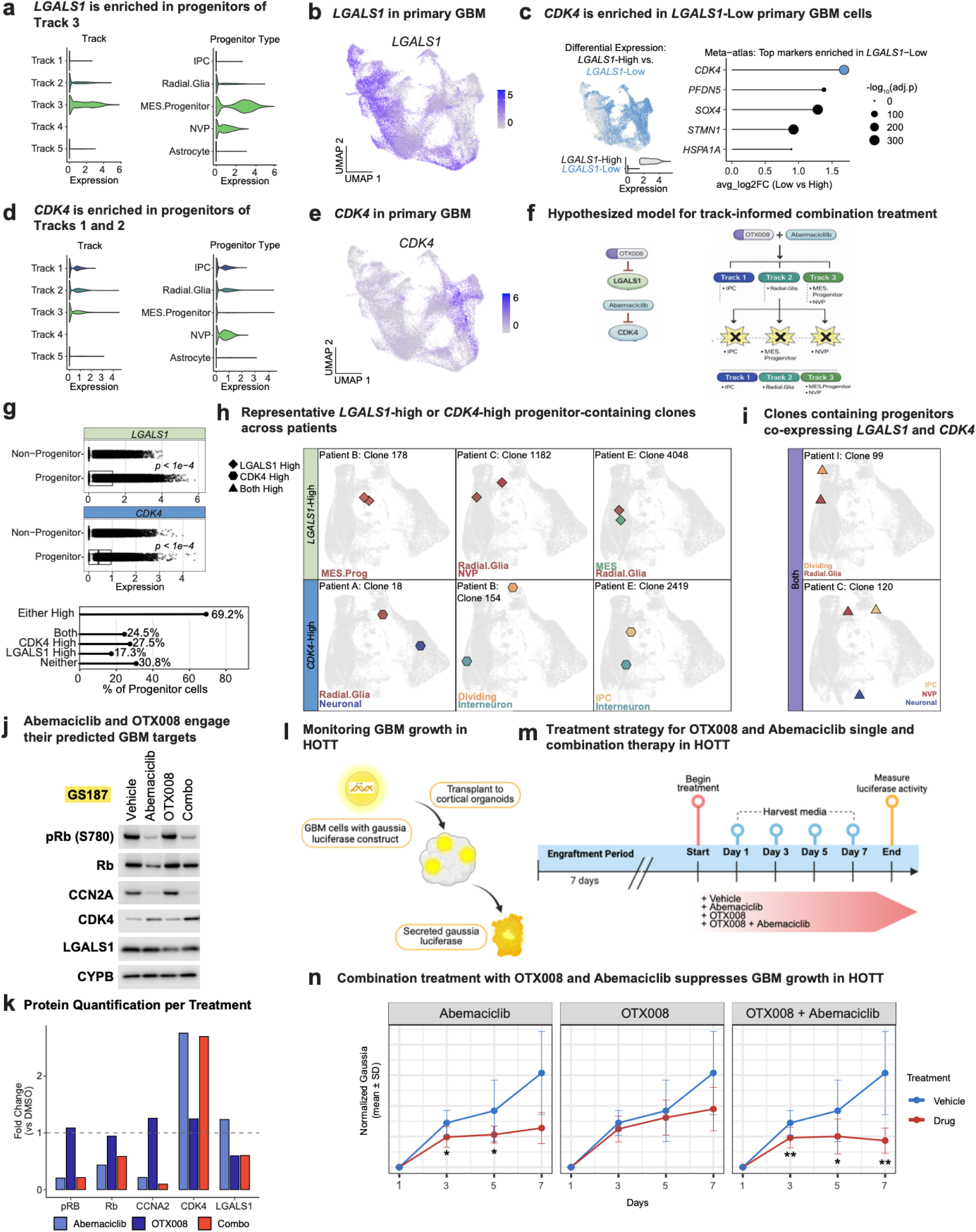
Track-informed combination targeting of progenitor populations suppresses GBM growth a) *LGALS1* expression across progenitor Tracks (left) and progenitor cell types (right) in the CellTag lineage-tracing dataset. b) FeaturePlot of *LGALS1* expression in the GBM meta-atlas of primary tumor cells. c) Primary GBM meta-atlas cells were binned into *LGALS1*-High (grey) and *LGALS1*-Low (light blue) based on *LGALS1* expression (STable6, Methods): cells were split by the median *LGALS1* expression (High ≥ median, Low < median). Violin plot shows *LGALS1* expression in each bin. Differential expression was then performed Low vs High (so positive avg_log2FC indicates genes enriched in *LGALS1*-Low cells). Right, lollipop plot shows the top differentially expressed genes enriched in LGALS1-Low cells, highlighting CDK4 (dot size, −log10 adjusted P value; x-axis, avg_log2FC Low vs High). d) *CDK4* expression across progenitor Tracks (left) and progenitor cell types (right) in the CellTag lineage-tracing dataset. e) FeaturePlot of *CDK4* expression in the GBM meta-atlas of primary tumor cells. f) The combination of *LGALS1* inhibition through OTX008 with *CDK4* inhibition through Abemaciclib is predicted to target Track 3 progenitors (MES.Progenitor, NVP) and complementary Track 1-2 progenitors (IPC, Radial Glia). g) *LGALS1* and *CDK4* expression stratifies progenitor cells into complementary high-expression groups. (Top) Boxplots show *LGALS1* and *CDK4* expression in the CellTagging dataset comparing Progenitor vs Non-Progenitor cells. P-values are from two-sided Wilcoxon tests. (Bottom) Progenitor cells were additionally binned by global median splits of LGALS1 and CDK4 across all cells to define four mutually exclusive categories (Neither, LGALS1_high, CDK4_high, Both) and an OR category (Either_High = LGALS1_high OR CDK4_high). Lollipop plot reports the percent of progenitor cells in each category. h) Representative clones satisfying LGALS1_high or CDK4_high treatment categories across patients, as defined in panel g. Clone member cells are overlaid on the global UMAP (grey) and indicated by colored shapes, with each shape representing an individual cell within the clone. i) Representative clones multiple patients containing cells classified as Both (LGALS1_high and CDK4_high), as defined in panel g. j) Representative immunoblot of the patient-derived gliomasphere line GS187 treated for 7 days with Vehicle, Abemaciclib (CDK4/6 inhibitor), OTX008 (LGALS1 inhibitor), or the combination. Abemaciclib treatment reduced phosphorylation of RB at S780 (pRb) with corresponding effects on total RB, consistent with on-target CDK4/6 pathway inhibition. CCN2A and CDK4 levels are shown as additional markers in the CDK4/6 axis. OTX008 treatment targets LGALS1 and CYPB serves as a loading control. k) Quantification of immunoblot signals shown in panel 4j from GS187 gliomaspheres. Protein levels for pRB (S780), RB, CCN2A, CDK4, and LGALS1 were normalized to the loading control (CYPB) and expressed as fold change relative to Vehicle (dashed line at 1). l) Patient-derived gliomaspheres expressing a Gaussia luciferase construct are transplanted onto human cortical organoids, where tumor cells engraft and secrete luciferase into the surrounding media. Secreted luciferase activity in conditioned media provides a noninvasive readout of tumor cell burden over time. m) Schematic timeline of the treatment regimen used for growth assays in the HOTT system. Following a 7-day engraftment period, organoids transplanted with patient-derived gliomaspheres were treated with Vehicle, Abemaciclib, OTX008, or the combination of OTX008 and Abemaciclib. Conditioned media was collected every 48 hours for measurement of secreted Gaussia luciferase activity, with luminescence quantified to generate tumor growth curves. n) Patient-derived gliomasphere lines GS005 and GS025 (n = 2 independent lines) expressing Gaussia luciferase were transplanted onto human cortical organoids (n = 3 independent transplants per condition) and treated according to the scheme in Fig. 4m. Conditioned media were collected every 48 hours and luciferase activity was normalized to the corresponding day 1 measurement for each transplant. Data are shown as mean ± s.d. across independent transplants. Statistical comparisons were performed between drug-treated and vehicle conditions at each time point.

Based on progenitor score, the cell types neatly separated into two significantly distinct groups: one containing NVP, Astrocyte, Dividing, Radial.Glia, MES.Progenitor and IPC with the other group containing Oligodendrocyte, Immune.Reactive, Newborn.Neuron, MES, Neuronal, and Hypoxic cells (Fig. 3b, SFig. 4b). Surprisingly, the cell population annotated as Astrocyte (due to its expression of *AQP4* and *GFAP* highly expressed the progenitor module. Upon further investigation, this Astrocyte-like population indeed co-expressed canonical progenitor markers such as *SOX2* and *NES* (SFig. 2d). When examining its clone relationships, this cell type was closely related to interneurons (SFig. 2c, STable4).

Although clonal relationships do not allow us to infer directionality of state transitions, we hypothesized that the progenitors we identify transcriptionally can self-renew and would have clonal relationships with more differentiated populations, reminiscent of progenitor-mature axes described in the literature. To establish that progenitor identities defined by CellTag lineage tracing reflect biologically meaningful and transferable states, we validated these annotations across independent datasets and modalities. To assess whether these progenitor programs generalize beyond the lineage tracing system, we projected them onto a published GBM Visium spatial transcriptomics dataset using reference-based label transfer (Methods). CellTag-defined progenitor identities mapped to spatially coherent domains within individual tumors and were associated with concordant expression of progenitor marker genes (SFig. 2d, 4c). Extending this analysis across six spatially profiled tumors demonstrated that these progenitor-associated transcriptional signatures reproducibly align with specific cell-type groupings across patients, supporting the broader relevance of CellTag-derived progenitor states (SFig. 4d). Together, these analyses establish that progenitor populations identified by lineage tracing correspond to conserved transcriptional and spatial programs in human GBM, enabling systematic interrogation of their clonal behaviors across tumors.

To understand how distinct progenitor populations contribute to tumor growth, we examined their relative participation in clonal relationships across patients. We noted that all progenitor populations were involved in clonal relationships in at least a subset of patients (Fig. 3c, SFig 4e). Radial glia contribute nearly half of progenitor clones to all patient samples interrogated in our HOTT system. The remaining progenitors, however, contribute variably to GBM clones and exhibit patient-specific functional relationships (Fig. 3c-h, SFig. 4e).

Although radial glia and dividing cells were the most abundant progenitor populations across tumors, the relative contributions of progenitors varied substantially between patients, with IPCs, NVPs, astrocyte-like progenitors, and MES.Progenitors each comprising sizeable fractions of progenitor-containing clones in specific tumors (Fig. 3c). An abundance-weighted co-occurrence analysis across patients further showed that progenitor types preferentially co-occur with other progenitors within the same clone, but with marked asymmetry in their interaction strengths (Fig. 3d). Radial glia exhibited the strongest and broadest co-occurrence with other progenitor populations, whereas IPCs, NVPs, astrocyte-like progenitors, and mesenchymal progenitors displayed more selective pairing patterns. This heterogeneity prompted us to examine how progenitor composition and co-occurrence patterns vary across tumors and in independent datasets.

Consistent with their annotation as progenitors, all progenitors had clonal relationships with their own cell type, suggesting an ability to self-renew. Radial glia were able to give rise to all other tumor progenitor types, while IPC, NVP, Astrocyte, and MES.Progenitor cells had more restricted relationships (Fig. 3d,g, SFig. 4e). Each progenitor had a distinct footprint of progenitor and non-progenitor types that they were most commonly related to (Fig. 3h, SFig. 4e). While progenitors contribute variably, a key observation is that no progenitor makes every cell type in the tumor, and both within and cross-track relationships of each progenitor are observed (Fig. 3i). Notably, although radial glia gave rise to the other progenitor populations, they did not directly generate all differentiated tumor states, suggesting that targeting radial glia alone would leave other progenitor compartments intact and capable of sustaining tumor heterogeneity.

A large proportion of clones in our dataset contain at least 1 progenitor and at least 1 mature cell (Fig. 3j). Separating the cells across this axis allows us to see the flow of putative differentiation by cell type and visualize the cross track interactions (Fig. 3k). We observe that some progenitors are more restricted and mostly give rise to cells within their same track, while some are more promiscuous, exhibiting higher proportions of their relationships as cross-track (Fig. 3k-l). Of note, even the highly promiscuous track 3 still has features of fate restriction, and does not contain all cell types within individual clonal relationships; yet, the progenitors from this track, namely NVP and MES.Progenitor, were identified as the ones most often containing cross-track relationships.

### Integrating lineage-informed progenitor states to guide combinatorial therapeutic targeting

Our lineage-tracing analysis revealed that progenitor populations differ in both hierarchical position and propensity for cross-track differentiation. Radial glia-like cells occupy a dominant position within the hierarchy, while NVP and MES.Progenitor populations contribute disproportionately to cross-track clonal relationships. We therefore asked whether these lineage-informed distinctions could be leveraged to nominate perturbations targeting functionally complementary progenitor compartments. To identify candidate targets, we filtered genes from the progenitor expression signature for known drug-gene interactions using The Drug-Gene Interaction Database (DGIdb)^34^ and ranked candidates by selectivity for tumor versus normal brain (SFig. 5a-b, Methods). This approach prioritized genes that were both enriched in progenitor populations within the CellTag dataset and selectively expressed in malignant cells.

**Figure 5.**
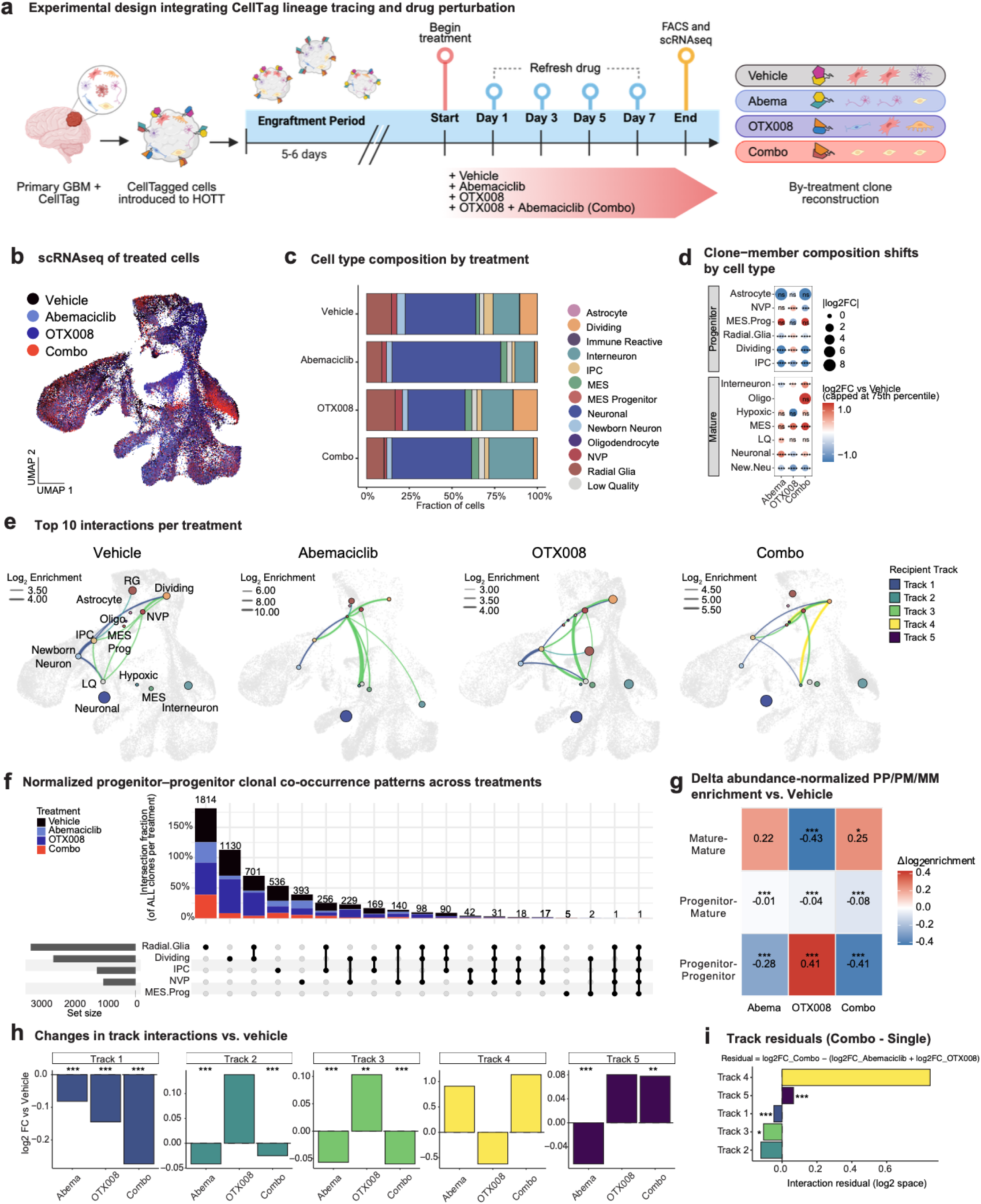
Track-informed combination treatment predictably remodels clonal interaction architecture a) Schematic of the track-informed CellTag lineage tracing and drug perturbation workflow in HOTT. Primary IDH1^WT^ GBM cells from (n = 3) patient tumors are dissociated and transduced with a high-complexity CellTag lentiviral barcode library encoding combinatorial DNA barcodes and GFP. Barcoded tumor cells are directly transplanted onto human cortical organoids and allowed to engraft and invade toward the organoid center over a 5-7 day period. Following engraftment, organoids are separated and treated with vehicle (DMSO), abemaciclib (1 µM), OTX008 (2.5 µM), or the combination of abemaciclib and OTX008 for 7 days, with drugs refreshed at each media change. At the experimental endpoint, organoids are dissociated and GFP⁺ CellTagged tumor cells are isolated by fluorescence-activated cell sorting (FACS) for single-cell RNA sequencing (scRNA-seq). Lineage relationships are reconstructed by identifying shared CellTag barcode combinations across cells, enabling analysis of treatment-induced changes in clonal architecture. b) UMAP visualization of scRNA-seq profiles from CellTag-labeled GBM cells following treatment in the HOTT (n = 98,474 cells). Cells are colored by treatment condition. c) Fraction of cells assigned to each transcriptional cell type following reference-based mapping from the full CellTag lineage-tracing dataset. d) Clone-member composition shifts (n = 9,138 clones) by cell type following drug treatment. Dot plot shows the log₂ fold change in clone-member representation for each cell type relative to vehicle, computed from unweighted clone-member proportions. For each treatment, the proportion of clone-member cells assigned to each cell type was calculated and normalized to the corresponding vehicle proportion (log₂FC). Dot size represents the magnitude of the log₂ fold change (|log₂FC|), and color denotes directionality (red, enrichment; blue, depletion), with values capped at the 75th percentile. Statistical significance was assessed at the clone level by comparing the distribution of clone-level cell type proportions between each treatment and Vehicle using two-sided unpaired Wilcoxon rank-sum tests, with p-values corrected across all cell type × treatment comparisons using the Benjamini–Hochberg method (**** FDR < 1×10⁻⁴, *** FDR < 1×10⁻³, ** FDR < 1×10⁻², * FDR < 0.05; ns, not significant). e) Top enriched clonal co-occurrence interactions per treatment. For each treatment condition (vehicle, abemaciclib, OTX008, combination), edges represent cell type pairs that co-occur within inferred clones more frequently than expected by chance (log₂ observed/expected), computed from clone × cell type presence matrices. Interactions were filtered to retain pairs with log₂ enrichment ≥ 0.5 and the top 10 strongest enriched pairs per treatment were plotted. Edge thickness denotes log₂ enrichment, and edges are colored by the Track assignment of the recipient cell type. f) UpSet plot of progenitor-progenitor clonal co-occurrence patterns across treatments. Columns denote the top 20 progenitor-progenitor cell type combinations observed within inferred clones; filled circles indicate presence of each progenitor cell type in the combination. The upper bar plot reports intersection size as the fraction of all clones within each treatment exhibiting the corresponding combination, with bars stacked by treatment (vehicle, abemaciclib, OTX008, combination). Fractions are normalized independently within each treatment using the total number of clones in that treatment as the denominator; therefore, stacked bar heights can exceed 100% because per-treatment fractions are summed across conditions. Left bars indicate progenitor set sizes. g) Delta abundance-normalized log₂ enrichment of progenitor-progenitor (PP), progenitor-mature (PM), and mature-mature (MM) cell-cell pairings for each treatment relative to Vehicle. Enrichment values reflect pooled clone-derived cell-cell interactions normalized to overall progenitor abundance. Significance was assessed at the clone level using two-sided Wilcoxon tests with Benjamini-Hochberg correction (* q < 0.05, ** q < 0.01, *** q < 0.001). h) Weighted clone-member Track composition changes relative to Vehicle. Bars show weighted log₂ fold change in Track representation among clone-member cells for each treatment versus Vehicle, with weights defined as 1 / p(Track | Treatment) and baseline Track frequencies computed from all cells. Significance was assessed using clone-level replication (Wilcoxon signed-rank test against 0) with FDR correction (* FDR < 0.05, ** FDR < 0.01, *** FDR < 0.001). i) Clone-level Track interaction residuals for combination treatment. Bars show deviation of the observed Combo effect from the additive expectation of Abemaciclib and OTX008, defined as log₂FC_Combo − (log₂FC_Abemaciclib + log₂FC_OTX008). Residuals were tested against 0 using clone-level replication (Wilcoxon signed-rank test) with FDR correction (* FDR < 0.05, ** FDR < 0.01, *** FDR < 0.001).

Among these candidates, *LGALS1* emerged as a prominent target with strong enrichment in Track 3 progenitors, including MES.Progenitor and NVP populations (Fig. 4a). Galectin-1 (*LGALS1*) has been shown to regulate neural and tumor progenitor cell states, and its genetic or pharmacological inhibition reduces tumor growth and modulates tumor-associated immune responses in preclinical GBM models^35–38^. *LGALS1* also predicts patient survival in proneural GBM, with *LGALS1*-high patients exhibiting lower overall survival (SFig. 5c). Consistent with these prior observations, *LGALS1* expression in our data was restricted to a subset of malignant cells in the GBM primary tumor meta-atlas and was not uniformly expressed across tumor populations (Fig. 4b). Because *LGALS1* expression was enriched in Track 3 progenitors, and our lineage analysis demonstrated that other progenitor populations also contribute substantially to tumor maintenance, we reasoned that targeting *LGALS1*-high compartments alone would leave *LGALS1*-low progenitors capable of sustaining tumor growth through compensatory expansion. This led us to search for a complementary target enriched in the LGALS1-low progenitor populations. Stratifying primary tumor cells by median *LGALS1* expression revealed a transcriptional program associated with *LGALS1*-low cells, in which *CDK4* was among the most significantly enriched genes (Fig. 4c, SFig. 5d, Methods). *CDK4* expression exhibited a complementary pattern to *LGALS1*, showing enrichment in Track 1-2 progenitors, including radial glia and IPC populations, while being comparatively reduced in Track 3 progenitors (Fig. 4d). Feature-based projection of *CDK4* expression in the GBM meta-atlas confirmed that CDK4 marks a distinct subset of tumor cells that only partially overlaps with *LGALS1*-expressing populations (Fig. 4e).

This complementary expression pattern is notable in light of prior clinical experience with CDK4/6 inhibition in GBM. While dysregulation of the CDK4/6-RB axis is a defining feature of GBM, and CDK4/6 inhibitors have demonstrated tumor penetration and on-target pathway inhibition, these agents have shown limited antitumor activity as monotherapies in both recurrent and newly diagnosed disease^39,40^. Our lineage-informed expression analysis provides a functional framework that may help explain these outcomes, as *CDK4* expression is enriched in specific progenitor compartments while remaining absent from other progenitor programs that contribute to tumor maintenance and cross-track differentiation. Although abemaciclib preferentially targets CDK4, CDK6 expression was uniformly distributed across Tracks 1, 2, 3, and 5 (SFig. 5e).

Based on these observations, we hypothesized that dual *LGALS1* and *CDK4* inhibition would simultaneously target functionally complementary progenitor populations, constraining the compensatory dynamics that sustain tumor hierarchy. We therefore predicted that combination treatment would suppress tumor growth more effectively than monotherapy and reduce the clonal contributions of Tracks 1–3. To pharmacologically target these programs, we selected OTX008 as a small-molecule inhibitor of LGALS1 and abemaciclib as a clinically validated CDK4/6 inhibitor (Fig. 4f). To assess how these targets partition the progenitor compartment in a Track-agnostic manner, we compared *LGALS1* and *CDK4* expression between progenitor and non-progenitor cells in the CellTagging dataset. Both genes were significantly enriched in progenitor populations (Fig. 4g, top). Progenitor cells were then binned into four mutually exclusive categories based on global median expression of *LGALS1* and *CDK4*. This analysis revealed that the majority of progenitor cells expressed either *LGALS1* or *CDK4*, with a smaller subset co-expressing both genes (Fig. 4g, bottom).

We sought to investigate whether *LGALS1* and *CDK4* were also in clones related to their predicted function. Previous work characterizing both candidates has identified them as potential drivers of glioma stem cell renewal^35–38^, leading us to explore their distribution across clones in our dataset. Lineage-traced clones containing *LGALS1*-high or *CDK4*-high progenitor cells were observed across multiple patients and occupied distinct regions of transcriptional space (Fig. 4h). Clones containing progenitors co-expressing both *LGALS1* and *CDK4* were also detected, confirming that these genes indeed are broadly expressed within tumor clones (Fig. 4i). Together, these findings provide lineage-resolved, computational support for a combinatorial perturbation strategy targeting complementary progenitor programs.

To test whether this lineage-informed combination therapy outperformed single agents, we first confirmed pharmacologic engagement of intended targets in (n = 3) patient-derived gliomaspheres (Fig. 4j-k, SFig. 5g). In order to monitor tumor growth, we transplanted patient-derived gliomaspheres which express secreted Gaussia luciferase^41^ onto human cortical organoids. After 7 days of engraftment in HOTT, treatment with Vehicle, Abemaciclib alone, OTX008 alone, or OTX008+Abemaciclib (Combo) was initiated (Fig. 4l-m, Methods). While single-agent treatment with either abemaciclib or OTX008 produced only modest effects on tumor growth, the combination resulted in a significant and sustained reduction in tumor burden by day 7 (Fig. 4n), suggesting cooperative effects beyond those observed with either agent alone.

### Track-informed combination treatment decreases progenitor-progenitor clone interactions

Having identified druggable targets enriched in distinct progenitor populations with combinatorial impacts on tumor growth, we next sought to explore if our proof-of-concept framework of using lineage relationships to predict combinatorial therapeutic targeting would impact tumor lineages as predicted. We hypothesized that simultaneous perturbation of radial glia-enriched Track 2 and NVP- and MES.Progenitor-enriched Track 3 pathways would remodel clonal architecture.

To test this hypothesis, we integrated CellTagging with pharmacologic perturbation in the HOTT system. Primary GBM cells from (n = 3) tumors were dissociated, infected with the CellTag lentiviral library, and transplanted onto human cortical organoids. Following an engraftment period of 5-7 days during which tumor cells invaded into the organoid center, transplants were randomly assigned to treatment conditions: Vehicle (DMSO), abemaciclib, OTX008, or a combination of the two (Combo). All conditions received equivalent Vehicle volumes. Drugs were refreshed with each media change over a 7-day treatment period, after which transplants were dissociated and GFP+ CellTagged cells were isolated by FACS for scRNA-seq (Fig. 5a-b, Methods). Our resulting dataset consisted of 98,474 cells and 9,138 clones. Across all three tumors, similar proportions of cells met CellTag barcode quality thresholds and were assigned as clone members, indicating consistent lineage-tracing coverage suitable for comparative clonal analyses (SFig. 6a).

Cells were annotated using reference-based mapping from the full CellTag dataset (Fig. 1c, Methods), yielding high-quality transcriptomes across all treatment conditions (Fig. 5b). Reference-based mapping using the CellTag lineage dataset enabled consistent assignment of progenitor and differentiated GBM cell states across treated samples, providing a common annotation framework for downstream clonal and track-level analyses(Fig. 5c, SFig. 6c). Despite impacts on tumor growth (Fig. 4m), cell type proportions exhibited modest variation across treatments, and both cells and clones were uniformly represented across conditions (SFig. 6d). However, clone size distributions shifted between treatment conditions, consistent with differential effects on progenitor expansion versus differentiation (SFig. 6b). Examination of clone-member composition revealed treatment-specific remodeling of clonal relationships. Compared to Vehicle, Combo treatment increased the representation of mature cell types, particularly mesenchymal and neuronal cells within individual clones, while decreasing the prevalence of progenitor-progenitor clone partnerships (Fig. 5d). Specifically, clones containing NVP, dividing cells, and IPC progenitors were reduced in Combo-treated samples, whereas clones containing oligodendrocytes and mesenchymal cells were enriched. Relative to single agent treatment, the drug combination treatment preferentially enhanced Track 4 clonal relationships while suppressing Tracks 1-3, consistent with a rebalancing away from progenitor-dominated clonal modules and toward more differentiated endpoints (Fig. 5e, SFig. 6g).

To systematically quantify treatment-induced changes, we calculated normalized clonal co-occurrence frequencies for every pair of progenitor states across all four conditions (Fig. 5f). In Vehicle-treated samples, progenitor interactions followed the expected hierarchy, with radial glia, dividing cells, and IPCs forming the most frequent clonal partnerships. Abemaciclib or OTX008 alone modestly reduced select interactions, whereas the combination produced a coordinated, robust suppression of progenitor–progenitor co-occurrence across all major progenitor pairs (Fig. 5f–g, SFig. 6e). After normalizing to Vehicle and stratifying by maturation state, combination treatment significantly reduced both progenitor–progenitor and mature–mature clonal interactions, while modestly increasing progenitor–mature partnerships (Fig. 5g). Track-level clonal diversity was not significantly different from Vehicle (SFig. 6f). Together, these results suggest that dual targeting disrupts progenitor self-renewal without simply forcing terminal differentiation, instead shifting clonal relationships toward intermediate differentiation trajectories.

Finally, we examined treatment effects on individual Track interactions. Compared to Vehicle, combination treatment significantly reduced interactions within Tracks 1, 2, and 3, the predicted Tracks of their putative targets, while increasing interactions within Tracks 4 and 5 (Fig. 5h). This remodeling was specific to the Combo; neither monotherapy recapitulated the full shift in Track connectivity (Fig. 5e, h, i). To isolate the combinatorial effect, we calculated interaction residuals by subtracting the sum of single-agent effects from the combination treatment effect for each Track. Positive residuals in Tracks 4 and 5, coupled with negative residuals in Tracks 1-3, confirmed that the combination produced emergent effects beyond the additive contributions of abemaciclib and OTX008 alone (Fig. 5i). Combination treatment produced non-additive, Track-specific remodeling of clone-member composition, with significant depletion of Tracks 1–3 and relative enrichment of Tracks 4–5 compared to either single agent alone (SFig. 6g). Track-resolved enrichment of combo-specific progenitor markers revealed distinct pathway programs across Tracks 1-3, consistent with heterogeneous non-additive transcriptional responses to combination treatment (SFig. 6h).

Together, these results establish a proof-of-concept that lineage-informed combinatorial targeting can predictably reshape GBM clonal architecture. By simultaneously perturbing radial glia-enriched (*CDK4*) and track 3 progenitor-enriched (*LGALS1*) pathways, combination treatment disrupted the progenitor-progenitor interactions that sustain tumor hierarchies, shifting clone composition toward more differentiated states in a manner consistent with the predicted effects based on progenitor lineage relationships. This framework provides a generalizable strategy for rational design of hierarchy-informed therapeutic combinations in GBM.

## Discussion

A major limitation in GBM therapeutic development has been the absence of functional frameworks linking transcriptionally defined cell states to tumor maintenance and plasticity. The classical hierarchical cancer stem cell model in GBM proposes that stem-like phenotypes may represent invariant stem cell hierarchies^42^, while more recent models posit that tumor organization may reflect dynamic state transitions within a plastic network^43^. While these studies establish that GBM cells can interconvert between transcriptional states^2^, they do not define how lineage relationships constrain or organize these transitions in primary human tumors. Using direct-from-patient human GBM samples, we analyzed 10,880 clones and found reproducible progenitor clonal patterns across tumors, with both constrained and cross-boundary capabilities. This merges the existing paradigms in the field, and provides a new framework for understanding GBM lineage progression. Despite substantial inter- and intra-tumoral heterogeneity, tumor genetics and patient context biased progenitor abundance and clone frequencies, but did not fundamentally alter how progenitor classes behaved. Rather than a purely stochastic equilibrium of interchangeable states, our data support a model in which plasticity operates within reproducible clonal architecture.

A central insight from our analysis is that no single progenitor population is sufficient to generate all tumor cell types. Even radial glia, despite occupying a relatively upstream position within the hierarchy and generating multiple other progenitor populations, do not directly produce all differentiated tumor states, indicating that a therapeutic strategy targeting radial glia alone would leave other progenitor compartments intact and capable of sustaining tumor heterogeneity. This distributed organization contrasts with models in which individual stem-like populations are proposed to fully regenerate tumor heterogeneity^5,44^, and instead supports a framework in which multiple progenitor classes contribute non-redundant lineage output.

Notably, progenitor populations differed markedly in their ability to traverse organizational axes. While many progenitors were largely constrained within a single Track, others exhibited frequent cross-Track relationships, indicating greater plasticity. Among all progenitors examined, the neurovascular progenitor (NVP) showed the highest propensity to participate in cross-Track clonal relationships. This distinguishes NVP from other progenitors and suggests that it may function as a lineage-intermediate population capable of linking otherwise constrained compartments. While the present study establishes this behavior at the level of clonal organization, these findings motivate further focused investigation into the mechanisms and functional consequences of NVP-mediated cross-axis plasticity. Recent multi-omic analyses have highlighted that GBM cell states are organized within epigenetically regulated hierarchies in which specific transcriptional regulators constrain or permit defined transitions^6^, supporting the existence of state-specific regulatory architectures. Our identification of NVP as a progenitor population with enhanced cross-Track connectivity provides lineage evidence for such asymmetric transition potential within the tumor ecosystem.

The emergence of drug-tolerant persister populations following targeted therapy has been attributed to reversible chromatin remodeling and activation of developmental programs^5^, as well as to dynamic state-switching within barcoded GBM cultures^43^. These adaptive responses highlight the capacity of tumors to re-establish heterogeneity following perturbation. In parallel, combined lineage barcoding and single-cell transcriptomic analyses have demonstrated that resistance to RTK inhibition can emerge through the coexistence of reversible persister states and genetically amplified subclones, revealing a coordinated interplay between epigenetic tolerance and clonal selection^7^. However, these adaptive trajectories were largely defined in cultured gliomasphere systems and did not address how resistance mechanisms are organized within the native lineage architecture of primary human tumors.

Our findings extend this framework by demonstrating that compensatory responses are not randomly distributed across malignant populations, but are structured within compartments that shape the routes of therapeutic escape. Accordingly, perturbing individual progenitors is likely to be circumvented through compensatory activity by other progenitors occupying distinct clonal footprints, consistent with the observed failure of numerous monotherapies in the field, including abemaciclib^45^ used in our proof-of-concept studies. These results provide a rationale for coordinated targeting of functionally complementary progenitor populations to constrain multiple clonal axes simultaneously. Consistent with this strategy, we applied lineage information to nominate combinatorial perturbations and assess their effects on clonal architecture. In lineage tracing of drug-treated primary patient tumor cells, combination treatment redirected clonal output away from highly plastic trajectories and preferentially funneled tumor cells into lower-plasticity compartments, shifts associated with reduced tumor burden. In contrast, individual treatments produced partial and compensatory effects, particularly among lineage-crossing progenitors. Thus, beyond cataloguing clonal organization, our framework demonstrates that lineage architecture can be experimentally reshaped in a data-driven manner that correlates with functional outcome.

Together, this study establishes a functional map of progenitor behavior in human GBM, providing a lineage-informed view of tumor organization. By showing that lineage relationships self-organize into clonal groups and that specific progenitors exhibit the capacity to cross these axes, we develop a framework that provides a foundation for understanding GBM adaptability and for designing rational strategies to constrain it.

## Acknowledgements

We would like to thank the members of the Bhaduri Lab for their insightful advice and comments on the study. We would like to thank the Broad Stem Cell Research Center Flow Cytometry core for their help in isolating cells for this project, and Charina Julian (UCSF) and Suhua Feng (UCLA) for help with running sequencing. We would like to thank Sergey Mareninov and others at the Brain Tumor Translational Research Core at UCLA for enabling tumor sample acquisition. We would like to thank UCLA Technology Center for Genomics and Bioinformatics (TCGB) for their help with the whole exome sequencing library preparation and sequencing. We would like to thank Maximilian Haeussler (UCSC) for his help in compiling the genome browser. This study was generously funded by support to AB from: NIH NCI P50CA211015 including a Career Enhancement Program Award, The Sontag Foundation (Distinguished Scholar Award), V Scholar Award from The V Foundation, The Uncle Kory Foundation, The American Cancer Society (CSCC-Team-23-980262-01-CSCC), The Margaret Early Medical Research Trust, The Rose Hills Foundation and the Broad Stem Cell Research Center. Funding for EF was provided by the David Geffen Scholarship and the UCLA-Caltech Medical Scientist Training Program (T32GM152342). BW was supported by CIRM funding DISC0-14514, a collaborative grant with AB. BW was additionally supported by NIH/NIMH RF1MH132662 and NIH/NHGRI U24HG002371. DC was supported by National Cancer Institute Ruth L. Kirschstein National Research Service Award F30CA295084.

## Author Contributions

The study was conceptualized by E.F. and A.B. Experimental design was performed by E.F., A.B., and D.N. P.R.N. and B.W. assisted with bioinformatic pipeline development. E.F., D.J.A., S.B., M.X.L., W.G., R.L.K., H.A.T., C.T., D.C., C.V.A., T.P. performed experiments and informatics. Data interpretation was performed by E.F., and A.B. Primary tissue samples were provided by K.S.P. and L.M.L. Analysis was performed by E.F. The manuscript was written by E.F. and A.B. with input and edits from all authors.

## Data Availability

Our GBM meta-atlas and all original datasets generated in this study are browsable and downloadable in the UCSC Genome Browser: (https://gbm-celltag.cells.ucsc.edu), in which the data are also available as Seurat objects for download. New raw data collected from this study are deposited in dbGAP:phs003936.v1.p1.

## Methods

### Culture of human embryonic stem cells for cortical organoid derivation

The human embryonic stem cell line UCLA6 was maintained under feeder-free conditions as previously reported^46–48^. Cells were cultured on Matrigel-coated six-well plates in mTeSR Plus medium supplemented with 10% mTeSR Plus Supplement and 1× Penicillin-Streptomycin or Primocin. Cultures were fed every other day, and cells were passaged once cultures exceeded approximately 75% confluence. For routine passaging, ReLeSR was applied to cultures at room temperature for 1 min and subsequently aspirated. Plates were then incubated at 37 °C for 5 min to facilitate selective detachment. Cells were gently dissociated into small clusters and replated onto freshly Matrigel-coated plates at a split ratio of 1:4 or 1:6, depending on growth requirements. For cryopreservation, cells were processed following the standard passaging protocol up to the replating step. Instead of replating, cells were resuspended in 1 ml mFreSR per well, transferred to cryogenic vials, and placed at −80 °C for 24-48 h prior to transfer into liquid nitrogen for long-term storage.

### Dissociation of primary GBM tissue

Primary tumor specimens were obtained from patients undergoing surgical resection at Ronald Reagan Hospital at UCLA following informed consent and approval from the Institutional Review Board (IRB protocols #10-000655 and #21-000108). Fresh tumor tissue was mechanically minced into small pieces using a sterile scalpel and transferred to 5-ml microcentrifuge tubes containing 2.5 ml papain solution supplemented with 125 µl DNase. Samples were incubated at 37 °C for 45-60 min to enable enzymatic dissociation, with vigorous manual agitation for approximately 10 s every 5 min to promote tissue breakdown. Following enzymatic digestion, tissue was further dissociated by trituration and centrifuged at 300 × g for 5 min to pellet cells. The resulting cell suspension was passed through a 40 µm cell strainer to remove undissociated material and debris, after which cells were counted. To further eliminate debris and enrich for viable cells, suspensions were processed using an ovomucoid density gradient according to the manufacturer’s instructions for the Papain Tissue Dissociation Kit. Briefly, up to 2 × 10^7 tumor cells were resuspended in a mixture of ovomucoid inhibitor, DNase, and EBSS and gently layered onto an ovomucoid inhibitor cushion. Gradients were centrifuged at 70 × g for 6 min, after which the supernatant was removed. Tumor cell pellets were collected and resuspended in either culture medium or FACS buffer for downstream applications.

### CellTag lentiviral transduction and transplantation into the human organoid tumor transplant (HOTT) system

After fresh dissociation, tumor cells in single-cell suspension were transduced with lentivirus carrying the CellTag-V1 barcode library (Addgene), which has a reported complexity of 19,974 unique barcodes. Following dissociation, cells were resuspended in Sasai3 medium and exposed to CellTag lentivirus in the presence of polybrene (1:1000 dilution). Transductions were performed at 37 °C for 60 min with gentle rotation to promote uniform viral exposure. The volume of lentivirus added was calculated to achieve a target multiplicity of infection (MOI) of 4 using the following formula:

TU_total = (MOI × number of cells) / viral titer (TU µl⁻¹).

After transduction, cells were washed three times with warm PBS and Sasai3 medium to remove excess virus and polybrene. Cells were then resuspended in 20 µl of Sasai3 medium for transplantation. For organoid co-culture, 8-12-week-old human cortical organoids were transferred to the lid of a 10-cm tissue culture dish using wide-bore 1,000 µl pipette tips. Excess medium was carefully removed, and 10-15 µl of the tumor single-cell suspension was deposited directly onto the surface of each organoid using the hanging drop method. The lid was then inverted onto a 10-cm dish containing 10 ml of base culture medium to maintain humidity and prevent evaporation during incubation. Hanging drop co-cultures were maintained at 37 °C for 12-16 h. Following this incubation period, organoids were transferred to ultra-low-attachment six-well plates containing Sasai3 medium. Tumor cells typically surrounded the organoids and initiated inward migration within several days. Co-cultures were maintained for 12-18 d, with media exchanges performed three times per week, before samples were collected for downstream analyses.

### Dissociation of organoid-tumor transplants and FACS isolation of GFP-positive cells

Tumor-engrafted cortical organoids were transferred to 1.7-ml microcentrifuge tubes containing 1 ml papain solution supplemented with 50 µl DNase and incubated at 37 °C for 45-60 min. To promote efficient dissociation, samples were vigorously agitated by hand for approximately 10 s during the first 10 min of incubation, with this agitation repeated every 5 min thereafter. Following enzymatic digestion, tissue was further dissociated by trituration and centrifuged at 300 × g for 5 min. Cell pellets were resuspended in cold FACS buffer, passed through a 40 µm cell strainer to remove residual aggregates, and stained with DAPI to enable exclusion of non-viable cells. Live GFP-positive tumor cells (DAPI⁻/GFP⁺) were isolated by fluorescence-activated cell sorting and collected for downstream single-cell capture. Flow cytometry and sorting were performed on a Bio-Rad S3e or Aria III cell sorter. Sequential gating strategies were applied to exclude debris, remove doublets, and eliminate dead cells based on DAPI incorporation. Gates defining GFP positivity were established using batch-matched, non-transplanted control organoids to account for background autofluorescence.

### Single-cell capture and library preparation

Single-cell suspensions were processed using the 10x Genomics Chromium single-cell 3′ gene expression platform. Depending on the number of cells available at the time of capture, libraries were prepared using the Chromium v3.1, GEM-X, or High-Throughput 3′ workflows. All single-cell encapsulation and library preparation steps were performed on the 10x Chromium system according to the manufacturer’s protocols corresponding to each chemistry. Following library construction, sequencing was carried out on an Illumina NovaSeq 6000 or NovaSeq X platform using standard 10x Genomics-recommended configurations.

### Gene expression processing and definition of QC’d tumor cells

Single-cell RNA-sequencing (scRNA-seq) gene expression libraries were aligned and quantified using the 10x Genomics Cell Ranger pipeline (v8.0.0). Human samples were aligned to a modified GRCh38 reference genome in which CellTag UTR and GFP coding sequences (GFP.CDS) were incorporated as additional reference features, enabling detection of expressed lineage barcodes alongside endogenous transcripts. Cell-by-gene count matrices were generated using default Cell Ranger parameters. Downstream analysis was performed using the Seurat v5 R package^49^. Cells were initially filtered based on standard RNA quality metrics, retaining only cells with at least 500 detected genes, expression of genes detected in at least three cells, and less than 10% mitochondrial transcript content. These baseline QC thresholds were applied uniformly across samples. Recovered cell numbers were quantified per sample from filtered matrices and used in conjunction with the corresponding 10x chemistry to calibrate sample-specific multiplet expectations.

Following quality filtering, gene expression values were normalized using log normalization, variable features were identified, and data were scaled prior to dimensionality reduction. Principal component analysis (PCA) was performed on the scaled expression matrix. The number of principal components retained for downstream analyses was determined using a data-driven heuristic that compares the variance explained by each component to an empirical noise threshold. significant principal components (PCs) were selected based on the larger value between the square of the standard deviation of PCA scores (Seurat.Obj@reductions$pca@stdev^2) and the square root of the ratio of the number of genes to the number of cells plus one (sqrt(length(row.names(Seurat.Obj)) / length(colnames(Seurat.Obj))) + 1)^2^50^. When this criterion yielded fewer than a minimum informative set, a conservative upper bound was applied to ensure sufficient dimensional coverage. The selected principal components were subsequently used for neighborhood graph construction, clustering, and Uniform Manifold Approximation and Projection (UMAP) embedding. Doublets were identified using DoubletFinder^51^, with expected multiplet rates estimated on a per-sample basis from recovered cell counts and 10x chemistry specifications. Tumor versus non-tumor cell identity was determined using inferCNV^52^, comparing malignant cells to a fixed normal reference to identify large-scale copy number alterations consistent with tumor genomes. Only high-confidence tumor singlets passing RNA QC, doublet exclusion, and inferCNV-based tumor classification were retained for downstream analyses.

### Side library construction for barcode-CellTag linkage

To directly associate 10x cell barcodes with CellTag sequences, a targeted side library was generated from DNA isolated from the same 10x Chromium reactions used for gene expression profiling. Side library amplification employed primers designed to append Illumina-compatible adapters while capturing the CellTag region downstream of GFP. Primer sequences (5′→3′) were as follows:

Forward (prtlTruSeqR1_1): ACACTCTTTCCCTACACGACG

Reverse (TruSeqR2-eGFP-3′): GTGACTGGAGTTCAGACGTGTGCTCTTCCGATCTggcatggacgagctgtacaag

The reverse primer anneals downstream of the GFP coding sequence, ensuring amplification of the CellTag cassette linked to the 10x cell barcode. Library amplification was performed using a high-fidelity polymerase to append TruSeq-compatible adapters, followed by indexing PCR to introduce sample indices and full Illumina adapters. Size selection and purification were carried out using SPRI bead-based cleanup to enrich for the expected library fragment size.

### Construction of a custom side library reference

Side library reads were aligned to a custom CellTag reference constructed using Cell Ranger (v8.0.0) with cellranger mkref. The reference consisted of consensus CellTag sequences derived from the whitelisted CellTag library, concatenated with a constant backbone sequence encompassing flanking adapter and reporter elements: GGCATGGACGAGCTGTACAAGTAAACCGGT[CellTag]GAATTCGATGACAGGCGCAGCTTCCGAGGGATTTGAGATCCAGACATGATAAGATACATTGATGAGTTTGGACAAACCAAAACTAGAATGCAGTGAAAAAAATGCCTTATTTG

Consensus CellTag sequences were generated to collapse sequencing errors prior to reference construction. Finalized FASTA and annotation files were compiled into a Cell Ranger-compatible reference using cellranger mkref, and side library reads were aligned and quantified using standard Cell Ranger alignment workflows optimized for targeted barcode recovery.

### Side library data processing

CellTag clone calling was performed using sequencing data from a dedicated side library, processed independently from gene expression data. Clone inference was restricted to the QC-filtered tumor singlet population defined by the gene expression workflow to ensure that lineage relationships were inferred only among high-confidence malignant cells.

Within this QC-restricted set, cells were required to exhibit between 1 and 20 detected CellTags to be eligible for clone inference, excluding cells with insufficient barcode complexity or anomalously high barcode counts indicative of technical artifacts. CellTag counts were subsequently binarized, and the binarized matrices were used to calculate a Jaccard similarity matrix on a per-sample basis. Clonal groupings were defined by applying a similarity cutoff of 0.7, and connected components within the resulting similarity structure were designated as clones. CellTag similarity and clone-calling logic were adapted from previously described barcode-based lineage inference frameworks^26^.

Cells that did not meet eligibility criteria for clone inference were retained in the dataset but annotated as not contributing to clone calling. Clone identities were integrated into the gene expression metadata for each QC-passing cell, enabling clone-aware downstream analyses without modifying underlying gene expression quantification.

### Estimation of barcode collision rates

Barcode collision rates were estimated by Monte Carlo simulation using the empirically observed CellTag count distribution and a library complexity of 19,073 whitelisted barcodes. Jaccard similarity was evaluated only for cells with 2-20 detected CellTags, with tag counts sampled from the global empirical distribution conditioned on this range; CellTags were drawn with replacement, binarized to sets, and pairwise Jaccard similarity was computed, with Jaccard ≥0.7 defining a false-positive similarity event. The pairwise false-positive rate was estimated from 50 million simulated cell pairs, and dataset-level expectations were calculated using the number of clone-calling-eligible cells, assuming an Erdős-Rényi model for random similarity edges. No clone-calling constraints beyond the Jaccard threshold were applied.

### Reference-based label transfer using query-reference mapping

To compare and transfer cell type annotations across datasets, we performed reference-based label transfer using a query-reference mapping framework implemented in Seurat. In each analysis, one dataset was designated as the reference and contained predefined cell type annotations, while the second dataset was treated as the query and processed independently according to dataset-specific quality control and filtering criteria prior to mapping. Shared transcriptional structure between reference and query datasets was identified by computing anchors using canonical correlation analysis-based feature alignment. Query cells were then projected into the low-dimensional embedding of the reference dataset using the MapQuery function, and reference annotations were transferred to query cells based on nearest-neighbor relationships in the shared embedding space. This approach enabled consistent annotation of query datasets without reclustering or redefining cell identities and allowed direct assessment of annotation concordance across datasets. Label transfer was performed independently for each reference-query pairing used in the study.

### Cell type and state annotation of the CellTagging scRNA-seq dataset

Transcriptional cell types and states in the CellTagging dataset were annotated using an integrated strategy combining reference-based mapping, gene module scoring, and cluster marker analysis. First, cells were mapped to a curated glioblastoma (GBM) meta-atlas of primary IDH1-wild-type tumors using Seurat’s query-reference framework (MapQuery), enabling transfer of cell type and state labels based on nearest-neighbor relationships in shared embedding space^27,49^. In parallel, cells were scored for published GBM cell-state gene programs, including proneural-like and mesenchymal-like signatures, using curated gene sets from prior studies^2,18^. These scores were used to validate transferred labels and to identify hybrid or transitional states not fully resolved by reference mapping alone. To ensure consistency with dataset-intrinsic structure, we performed unsupervised clustering followed by cluster marker analysis using Seurat FindAllMarkers (only.pos = TRUE, min.pct = 0.25). Cluster-enriched genes were interpreted in the context of canonical GBM markers and human neurodevelopmental reference datasets, enabling alignment of malignant populations with neurodevelopmental analogs such as radial glia-like, intermediate progenitor-like, and neuronal-like states^53,54^. Final annotations were assigned by integrating concordant evidence across label transfer, module scoring, and cluster marker profiles.

### Clone-level cell type co-occurrence and enrichment analysis

Clone-level cell type co-occurrence and enrichment analysis. Clone-level relationships between transcriptionally defined cell types were quantified using CellTag-defined clonal identities. For each clone, a binary clone × cell type presence matrix was constructed, in which a cell type was scored as present if it appeared at least once in the clone, independent of cell number. Background cell type frequencies were calculated from all annotated cells in the dataset and used to define a null model of random cell type co-occurrence under independence. Observed co-occurrence between pairs of cell types was computed as the number of clones in which both types were present, and expected co-occurrence was estimated based on background frequencies and the total number of clones. Enrichment of co-occurrence was quantified as the log₂ ratio of observed to expected values, with a small constant added for numerical stability and diagonal entries excluded. The resulting enrichment matrix was hierarchically clustered to identify higher-order structure in clone-level relationships. Statistical significance of block structure was assessed using a global blockiness statistic, defined as the difference between mean absolute enrichment within dendrogram-defined blocks versus between blocks, with significance evaluated by permutation testing in which cell type labels were randomly reassigned across cells while preserving clone sizes. Based on hierarchical clustering, cell types were grouped into five major co-occurrence communities, termed Tracks, representing broad axes of clonal proximity rather than strict lineage relationships. A schematic overview of the co-occurrence enrichment calculation is shown, with representative numerical examples using real data provided in Supplementary Fig. 2e.

### Progenitor gene signature derivation and scoring

To derive a progenitor-associated transcriptional signature, the CellTagging dataset was randomly split into two non-overlapping halves. One half was used for signature discovery, excluding cells annotated as Dividing, and the other was reserved for independent validation. Differential expression analysis was performed in the discovery set comparing reannotated progenitor cells to non-progenitor malignant cells using Seurat FindAllMarkers (only.pos = TRUE, min.pct = 0.25, logfc.threshold = 0.25). For each progenitor-associated gene, a genescore was calculated to capture both specificity and enrichment, defined as the ratio of the fraction of progenitor cells expressing the gene to the fraction of non-progenitor cells expressing the gene (pct.1/pct.2), multiplied by the average log_2_ fold change. Genes were ranked by genescore, and the top 200 genes were selected to define the progenitor signature. The top 200 genes by genescore were used for progenitor module scoring in the held-out validation dataset, whereas the full, unfiltered progenitor gene set was retained for downstream druggability and target enrichment analyses.

### Progenitor-progenitor co-occurrence analysis and arc diagram visualization

To quantify relationships among progenitor populations, analyses were performed on a progenitor-only Seurat object containing cells annotated as IPC, Radial Glia, Dividing, MES.Progenitor, NVP, or Astrocyte. For each patient, the proportion of cells belonging to each progenitor subtype was calculated relative to the total progenitor compartment. Progenitor subtypes were considered present within a patient if they exceeded a minimum within-patient proportion threshold (2%). Pairwise progenitor co-occurrence was quantified in an abundance-weighted manner by computing, for each patient, the product of the proportions of each progenitor pair, and summing these values across patients to obtain a global co-occurrence weight. Node sizes in the arc diagram represent the average proportion of each progenitor subtype per patient, while edge widths correspond to the summed abundance-weighted co-occurrence across patients. To improve interpretability, weak edges below a minimum co-occurrence threshold were excluded, and progenitor subtypes were ordered based on network connectivity. The resulting arc diagram summarizes conserved and variable patterns of progenitor co-occurrence across tumors.

### Processing of the Nomura glioblastoma dataset

Single-cell RNA-seq data from the Nomura glioblastoma cohort were obtained and processed with attention to platform-specific and quality-control differences. Although the original study included data generated using both Smart-seq and 10x Genomics platforms, only the 10x Genomics dataset was used for downstream analyses, as reported cell counts and annotations in the original publication correspond to this subset. Initial quality control thresholds were applied to remove low-quality cells. While the original study retained cells with as few as 200 detected genes, we applied a more stringent filter requiring a minimum of 500 detected genes per cell, yielding approximately 600,000 high-quality cells. Cells were then stratified based on the original annotations into malignant, non-malignant, and unassigned (“NA”) categories, where NA indicated ambiguous copy number profiles. Cells annotated as non-malignant or NA were excluded from downstream analyses, resulting in a final set of 241,708 malignant cells. This number is modestly lower than reported in the original study, reflecting the additional quality-control filtering applied here. Malignant cells were subsequently annotated by reference-based label transfer using Seurat MapQuery, mapping query cells to the GBM meta-atlas to assign transcriptional cell type and state identities in a consistent framework.

### Gene ontology enrichment analysis

Gene ontology (GO) enrichment analysis was performed using the clusterProfiler package. Gene lists were tested for enrichment of Gene Ontology Biological Process (GO BP) terms using the enrichGO function with the human reference database (org.Hs.eg.db) and gene symbols as identifiers. Multiple testing correction was performed using the Benjamini-Hochberg method, and terms with adjusted P < 0.05 were considered significant.

### Visium spot annotation by RCTD

Visium Seurat objects from Tsyben et al.^55^ were downloaded and spot identities were assigned by RCTD (spacexr) using the GBM meta-atlas as the single-cell reference. An RCTD Reference was constructed from meta-atlas raw RNA counts with meta-atlas cell type labels, and Visium slices were combined by concatenating per-slice centroid coordinates and matching per-slice count layers prior to building the SpatialRNA query. RCTD was run in doublet mode (doublet_mode = “doublet”), and spot annotations were derived from the primary assignment (first_type), which was appended to the Visium Seurat object for downstream analyses and visualization.

### Identification and prioritization of druggable progenitor targets

To identify candidate therapeutic targets enriched in tumor progenitor populations, we focused on genes defining the unfiltered progenitor signature derived from the CellTag-based lineage tracing dataset. Progenitor signature genes were queried against the Drug-Gene Interaction Database (DGIdb) using its GraphQL API to identify genes with known drug interactions. A gene was considered druggable if DGIdb reported at least one interaction annotated with an inhibitor-related interaction type, including inhibitor, antagonist, or compound with inhibitory activity (case-insensitive match). Queries were performed in batches to retrieve all reported interactions, and only inhibitor-linked interactions were retained. This yielded a set of 326 progenitor-associated druggable genes.

To assess tumor selectivity, global mean expression levels for each druggable gene were computed across all malignant cells in a glioblastoma (GBM) meta-atlas and across all cells in a normal adult human brain meta-atlas^53^. Tumor versus normal enrichment was quantified by z-scoring mean expression values across genes within the GBM and normal brain datasets separately, then subtracting the z-scored normal brain expression from the z-scored GBM expression to generate a tumor-normal score. Positive values indicate relative enrichment in tumor compared with normal brain, whereas negative values indicate higher relative expression in normal tissue. Druggable progenitor-associated genes were prioritized by jointly considering their CellTagging progenitor genescore and their tumor-normal score, enabling identification of candidate targets that are both lineage-informed progenitor-specific and selectively enriched in glioblastoma relative to normal brain.

### LGALS1 High/Low stratification and differential expression analysis in the GBM meta-atlas

*LGALS1*-high and -low groups were defined within the primary GBM meta-atlas Seurat object using LGALS1 expression. Cells were assigned a binary LGALS1_bin label by a dataset-specific median split on LGALS1 expression.Differential expression between LGALS1-Low and LGALS1-High groups was performed using Seurat FindMarkers with Wilcoxon rank-sum testing (test.use = “wilcox”), min.pct = 0.5, and logfc.threshold = 0.25. For all analyses, ident.1 was set to LGALS1-Low and ident.2 to LGALS1-High; therefore, positive avg_log2FC values indicate genes enriched in LGALS1-Low cells. Genes were considered significant if they met the FindMarkers expression prevalence requirement (detected in at least 50% of cells in at least one group, as enforced by min.pct = 0.5) and had an adjusted p-value below 0.05. CDK4 enrichment within the LGALS1-Low group was evaluated directly from the FindMarkers output based on adjusted p-value significance and directionality (positive avg_log2FC, indicating higher expression in LGALS1-Low cells).

### Patient-derived GBM gliomaspheres and Gaussia luciferase labeling

All patient-derived glioblastoma tumor specimens were obtained following explicit informed consent under University of California, Los Angeles Institutional Review Board approval (IRB protocol #10-000655). Gliomasphere cultures were a generous gift from the Nathanson laboratory and were generated as described here. Fresh tumor tissue was subjected to combined mechanical and enzymatic dissociation using the Human Tumor Dissociation Kit (Miltenyi Biotec). Red blood cells were removed using red blood cell lysis buffer, followed by sequential depletion of CD45⁺ immune cells and myelinated cells using antibody-conjugated magnetic beads and two rounds of column-based magnetic separation. Primary GBM cells were established and maintained under gliomasphere culture conditions consisting of DMEM/F12 (Gibco) supplemented with B27 (Invitrogen), penicillin-streptomycin (Invitrogen), and GlutaMAX (Invitrogen), with additional supplementation of heparin (5 µg ml⁻¹; Sigma), epidermal growth factor (EGF; 20 ng ml⁻¹; Sigma), and fibroblast growth factor (FGF; 20 ng ml⁻¹; Sigma). Cultures were maintained at 37 °C in an atmosphere of 20% O₂ and 5% CO₂.

Gliomasphere cultures were transduced by spinfection with a lentiviral construct encoding a secreted Gaussia luciferase (sGluc) reporter (Prolume Ltd., pLenti_CMV_GLuc_T2A_eGFP plasmid). Following transduction, cells were dissociated into a single-cell suspension and transplanted into the human organoid tumor transplant (HOTT) system, enabling longitudinal, non-destructive monitoring of tumor burden through Gaussia luciferase secretion into conditioned media. All gliomasphere lines were routinely tested and confirmed negative for mycoplasma contamination using a commercially available assay (MycoAlert, Lonza). Cell lines were used at fewer than 15 passages and were authenticated by short tandem repeat (STR) profiling prior to experimental use.

### Immunoblotting

Cells were collected and lysed in RIPA buffer (Thermo Scientific, 89901) supplemented with Halt Protease and Phosphatase Inhibitor Cocktail (Thermo Scientific, 78440). Lysates were clarified by centrifugation at 14,000 × g for 15 min at 4°C. Protein samples were denatured in NuPAGE LDS Sample Buffer (Invitrogen, NP0007) and reduced with BondBreaker TCEP (Thermo Scientific, 77720), then separated by SDS-PAGE using 4–12% Bis-Tris gradient (Invitrogen, NP0329) or 12% continuous gels (Invitrogen, NP0343) and transferred to 0.2 μm PVDF membranes (Thermo Scientific, 88520). Blocking was performed in 5% non-fat milk in TBST. Membranes were incubated overnight with primary antibodies against pRb (S780; Cell Signaling Technology, 8180), Rb (Cell Signaling Technology, 9309), CCNA2 (Cell Signaling Technology, 4656), CDK4 (Cell Signaling Technology, 12790), CYPB (Cell Signaling Technology, 43603), and LGALS1 (Cell Signaling Technology, D608). Membranes were then incubated for 1 hour at room temperature with HRP-linked secondary antibodies: anti-rabbit IgG (Cell Signaling Technology, 7074) or anti-mouse IgG (Cell Signaling Technology, 7076). Membranes were developed using SuperSignal Pico PLUS (Thermo Scientific, 34580) or SuperSignal Femto (Thermo Scientific, 34094) chemiluminescent substrates and imaged on a LI-COR Odyssey FC system.

### Drug treatment and Gaussia luciferase-based tumor growth quantification in the HOTT system

Patient-derived gliomaspheres expressing secreted Gaussia luciferase were dissociated into single-cell suspensions and transplanted onto human cortical organoids using the hanging drop method. Following transplantation, organoid-tumor co-cultures were maintained for a 7-day engraftment period before initiation of drug treatment. After engraftment, organoid tumor transplants were treated with vehicle control (DMSO), abemaciclib (1 µM; MedChemExpress), OTX008 (2.5 µM; MedChemExpress), or the combination of abemaciclib and OTX008. All treatment conditions, including vehicle controls, received equivalent final concentrations of DMSO. Drug treatments were administered for 7 days, with compounds refreshed at each media change. Conditioned media were collected every 48 h for quantification of secreted Gaussia luciferase activity, which was used as a non-destructive measure of tumor burden over time. Organoid transplants were individually housed to enable longitudinal growth measurements. For each transplant, luciferase values were normalized to the corresponding day 1 measurement and summarized as mean ± s.d. across time. Statistical comparisons were performed between drug-treated and vehicle conditions at each time point.

### CellTag lentiviral transduction and transplantation into the human organoid tumor transplant (HOTT) system with drug treatment

Primary IDH1-wild-type glioblastoma specimens from three patients were dissociated into single-cell suspensions and transduced with a high-complexity CellTag lentiviral library encoding combinatorial DNA barcodes and GFP, as previously described. Immediately following transduction, barcoded tumor cells were transplanted onto human cortical organoids to establish human organoid tumor transplants (HOTT). Tumor cells were allowed to engraft and invade toward the organoid interior for 5-7 days, after which individual organoids were separated and randomly assigned to treatment conditions. Organoids were treated with vehicle control (DMSO), abemaciclib (1 µM, MedChemExpress), OTX008 (2.5 µM, MedChemExpress), or a combination of abemaciclib and OTX008 for 7 days, with compounds refreshed at each media change. At the experimental endpoint, organoids were dissociated to single cells, and GFP⁺ CellTagged tumor cells were isolated by fluorescence-activated cell sorting. Sorted cells were processed for single-cell RNA sequencing, and lineage relationships were reconstructed by identifying shared CellTag barcode combinations across cells.

### Track-stratified progenitor differential expression, gene set definition, and pathway enrichment

Differential expression analysis for Fig. 5g was performed on progenitor cells stratified by Track. For each Track, progenitor cells were subset and differential expression was computed between Vehicle and each treatment condition (Abemaciclib, OTX008, and Combo) using Seurat’s FindMarkers function, retaining only genes with increased expression in the treatment condition, detected in at least 25% of cells, and exceeding a log_2_ fold-change threshold of 0.25. Statistical significance was assessed using adjusted p-values, with an FDR threshold of 0.05. To identify combination-specific transcriptional responses, an outersect gene set was defined for each Track as genes significantly altered in the Combo versus Vehicle comparison that were not significant in either single-agent versus Vehicle comparison under the same thresholds. These Track-specific outersect gene sets were then subjected to pathway enrichment analysis using MSigDB Hallmark and Reactome gene set collections, with multiple-testing correction applied within each enrichment analysis

---

**Supplementary Figure 1.**
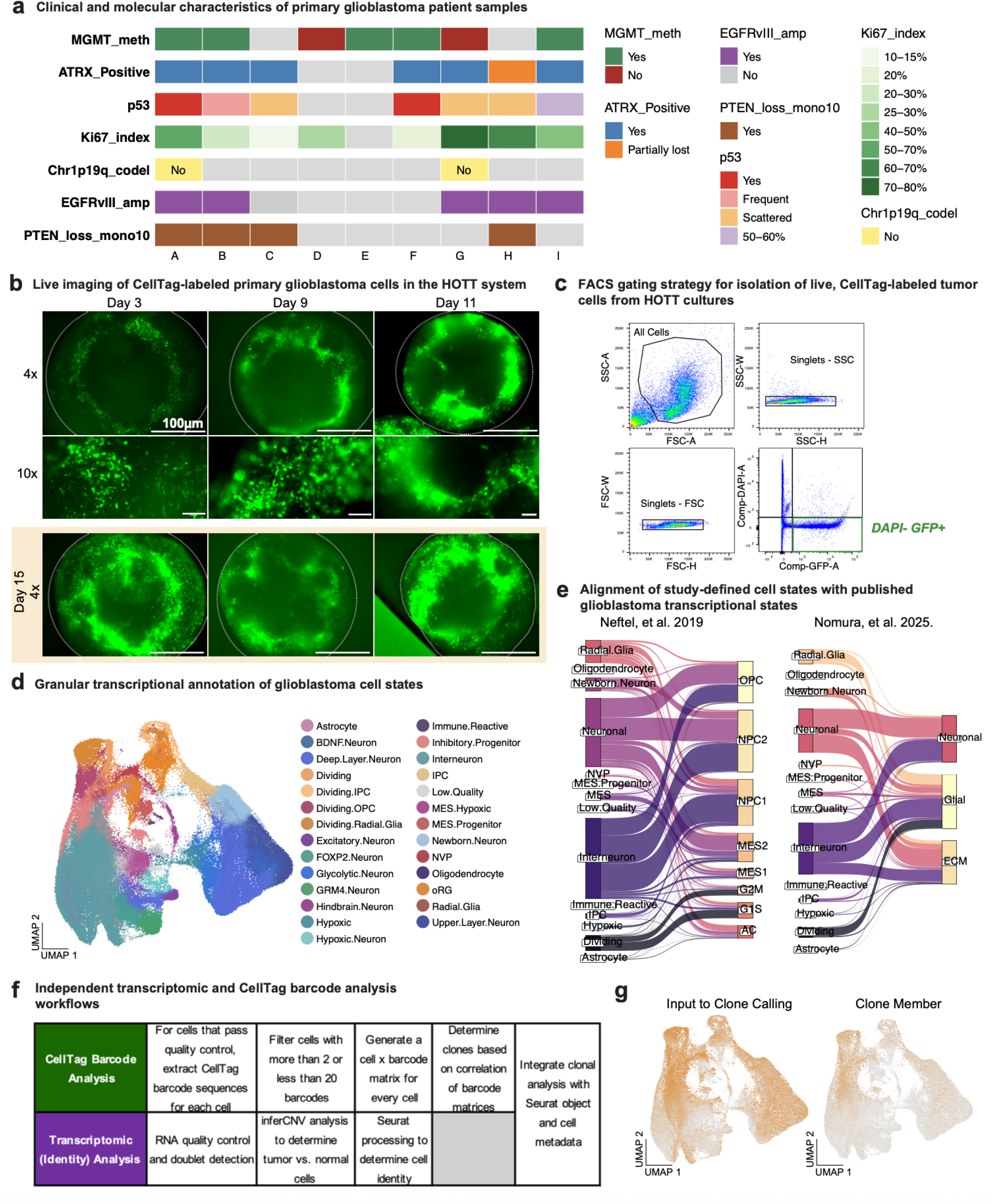
CellTag DNA barcoding and clonal analysis workflow for primary glioblastoma in the HOTT system a) j) Summary of clinical and molecular features for primary IDH1^WT^ GBM patient tumors used for CellTag-based lineage tracing experiments. Shown are key diagnostic and molecular annotations for each patient (A-I). Color coding indicates categorical or binned quantitative values as denoted in the legend. All tumors meet diagnostic criteria for primary GBM. Additional clinical and pathological information for each patient is provided in STable1. b) k) Live fluorescence imaging of CellTag-GFP-labeled tumor cells from Patient B following transplantation into human cortical organoids (HOTT system). Top and middle rows show representative images acquired at days 3, 9, and 11 post-transplantation at low (4×) and higher (10×) magnification, respectively. Organoid boundaries are indicated by white dotted outlines in the 4× images. Bottom row shows 4× images acquired at day 15, immediately prior to organoid harvest for downstream dissociation and single-cell RNA sequencing. Images illustrate progressive tumor cell expansion and infiltration within the organoid over time. Scale bars, 100 µm in all panels. c) l) Fluorescence-activated cell sorting (FACS) gating strategy used to isolate live tumor cells from dissociated human organoid tumor transplant (HOTT) cultures. Cells were first gated based on forward scatter (FSC-A) and side scatter (SSC-A) to exclude debris, followed by sequential singlet gating using SSC width versus height (SSC-W vs. SSC-H) and FSC width versus height (FSC-W vs. FSC-H). Live tumor cells were then identified as GFP-positive and DAPI-negative (GFP⁺ DAPI⁻), enabling separation of CellTag-labeled malignant cells from dead cells and unlabeled organoid-derived cells prior to single-cell RNA sequencing. d) m) UMAP visualization of the same 235,155 quality-controlled malignant GBM cells shown in Figure 1c, annotated at increased granularity to capture transcriptionally distinct subpopulations within major cell states. Cell identities were assigned using the same integrated annotation framework as in Figure 1c but allowing for additional subtyping where transcriptional resolution supported further distinction. These granular annotations were used for exploratory analyses and validation, while higher-level cell state groupings were applied for main figure analyses unless otherwise noted. e) n) Comparison of cell state annotations in panel 1c with previously published GBM transcriptional classification frameworks. Sankey diagrams depict the correspondence between study-defined high-level cell types (left) and cell states defined by Neftel et al., 2019^2^ (left panel) or Nomura et al., 2025^32^ (right panel), based on per-cell annotation overlap. Flow widths represent the number of cells assigned to each pairing. This analysis demonstrates that the cell states defined in this study largely align with major published GBM groupings while providing additional granularity. f) o) Following single-cell quality control, transcriptomic analyses were used to identify malignant tumor cells and assign cell states, while CellTag barcode extraction and clone identification were performed exclusively on quality-controlled malignant cells. Clonal information was subsequently integrated with transcriptomic annotations for downstream analyses. g) p) UMAP projections highlighting cells eligible for clone calling (left; orange) and cells assigned as clone members (right; orange). Eligibility reflects stringent CellTag barcode and quality filtering, and clone membership denotes cells assigned to inferred clones based on barcode similarity.

**Supplementary Figure 2.**
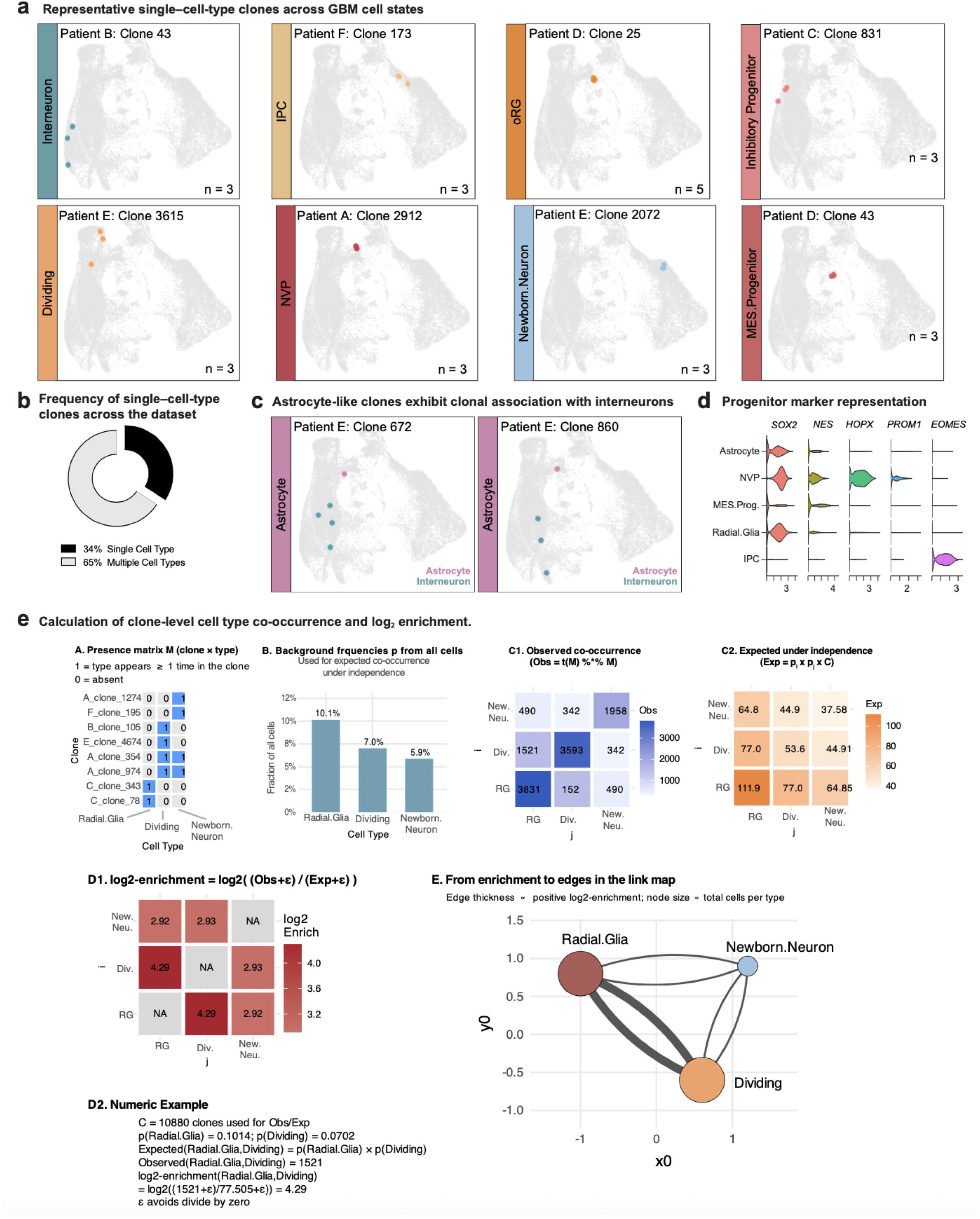
Single-cell-type clones and quantitative inference of clonal cell state co-occurrence a) Representative examples of single-cell-type clones identified by CellTag barcode similarity across multiple patients. Each panel shows one clone (colored points) projected onto the global UMAP of malignant GBM cells (gray). Colored dots represent individual cells belonging to the indicated clone and are colored by their annotated cell type, shown in the vertical label bar. Clone size (n) is indicated for each example. b) Proportion of inferred clones composed of a single transcriptional cell type versus multiple cell types c) Representative examples of multi-cell-type clones dominated by astrocyte-like progenitor identities. Although classified as astrocyte-like, these clones reproducibly include interneuron cells, supporting functional clonal relationships between astrocytic progenitors and interneuron fates. d) Violin plots showing expression of canonical neurodevelopmental progenitor marker genes across CellTag-defined progenitor populations e) Schematic overview of the analytical framework used to quantify enrichment of cell type co-occurrence within clones. (A) A binary presence matrix M encodes whether each cell type is present (≥1 cell) in each clone. (B) Background cell type frequencies (pᵢ) are calculated from all cells in the dataset and used to define the expected co-occurrence under independence. (C1) Observed co-occurrence is computed as t(M) × M, yielding the number of clones in which each pair of cell types co-occurs. (C2) Expected co-occurrence is calculated as pᵢ × po × C, where C is the total number of clones. (D1) Log₂ enrichment is computed as log₂((Observed+ε)/(Expected+ε)), with diagonal entries set to NA; ε is a small constant to avoid division by zero. (D2) A worked numeric example using real dataset values illustrates the calculation. (E) Enriched co-occurrence relationships are represented as edges in link maps, with edge thickness proportional to positive log₂ enrichment and node size proportional to total cell counts per type. See Methods.

**Supplementary Figure 3.**
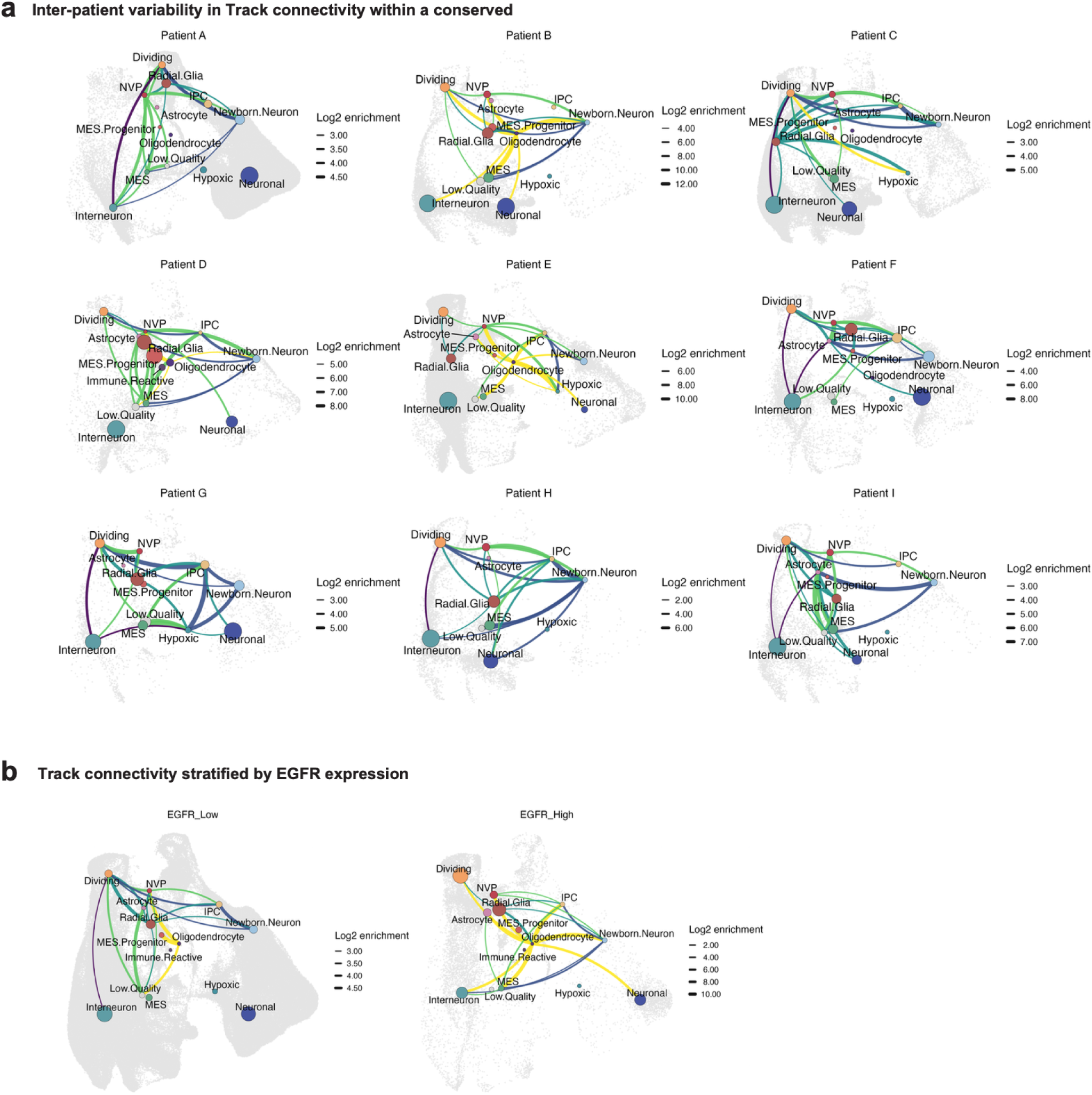
Conservation and variability of clonal Track structure across patients and genetic contexts a) Per-patient connectivity maps showing enriched clone-level co-occurrence relationships among GBM cell types across nine primary tumors. Nodes represent cell types positioned at the median UMAP coordinates of all cells of that type within each patient, with node size proportional to the total number of cells. Edges indicate significantly enriched clonal co-occurrence between cell type pairs (log₂ observed/expected ≥ 0.25), computed using patient-specific cell type abundances as the null expectation. Edge thickness reflects the magnitude of log₂ enrichment, and edges are colored by the developmental Track assignment of the recipient cell type. For each patient, the top 20 strongest enriched relationships are shown. b) Connectivity maps of enriched clonal relationships stratified by *EGFR* expression status, comparing *EGFR*-low and *EGFR*-high tumors. *EGFR* expression was quantified per cell using normalized transcript counts, and tumors were classified as *EGFR*-high or *EGFR*-low based on whether their median *EGFR* expression across malignant cells fell above or below the cohort-wide median. Node placement, sizing, and edge definitions are as in Fig. 3a.

**Supplementary Figure 4.**
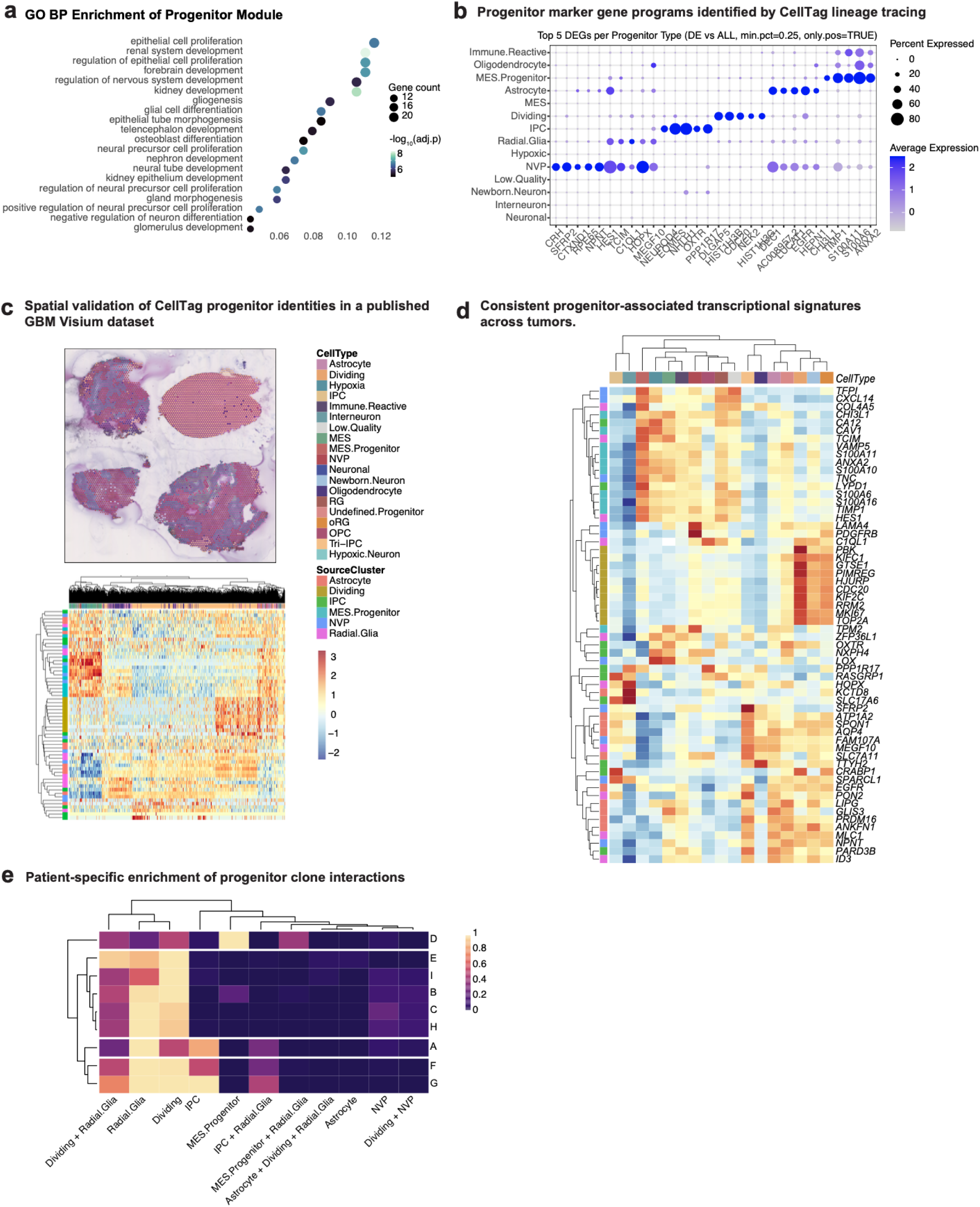
CellTag-defined progenitor states correspond to conserved transcriptional and spatial programs a) Gene ontology (GO) Biological Process enrichment analysis of the top 200 genes defining the progenitor transcriptional signature derived from the CellTagging dataset. Enrichment was performed using clusterProfiler as described in Methods. Each dot represents a significantly enriched GO term, plotted by gene ratio (x-axis) and term identity (y-axis). Dot size corresponds to the number of progenitor signature genes contributing to each term, and color indicates statistical significance expressed as -log₁₀(adjusted P value). Enriched terms highlight developmental, proliferative, and neural progenitor-associated programs consistent with progenitor cell identity. b) Dot plot showing the top differentially expressed genes for each progenitor population identified in the CellTag lineage tracing dataset. These markers define distinct transcriptional programs associated with individual progenitor identities used throughout downstream validation analyses. c) Representative Visium section (AT200_FO2_ii) annotated using RCTD-based label transfer from the GBM meta-atlas. Spots are colored by transferred cell-type labels, and expression of CellTag-defined progenitor marker genes is shown below, grouped by their source progenitor cluster. This analysis demonstrates that progenitor identities inferred from lineage tracing map onto spatially coherent transcriptional domains in an independent tumor sample (Methods). d) Average expression of CellTag progenitor marker genes across RCTD-annotated cell-type groups, aggregated over six Visium tumor sections. Rows represent marker genes grouped by progenitor source cluster, and columns represent annotated cell types. Hierarchical clustering reveals that CellTag-defined progenitor programs align with reproducible cell-type groupings across tumors, supporting their broader relevance beyond the original lineage tracing dataset (Methods). e) Most frequent progenitor-containing clone interaction patterns across patient tumors. Each interaction represents the combination of high-level cell types co-occurring within individual clones that contain at least one progenitor cell. For each patient, interaction frequencies were normalized by the total number of cells recovered from that patient to account for differences in sampling depth. Rows denote interaction labels (single or multi-cell-type combinations), and columns denote patients. Values are scaled within each patient (0-1) to highlight relative enrichment, with darker colors indicating interactions that are more prevalent for a given patient.

**Supplementary Figure 5.**
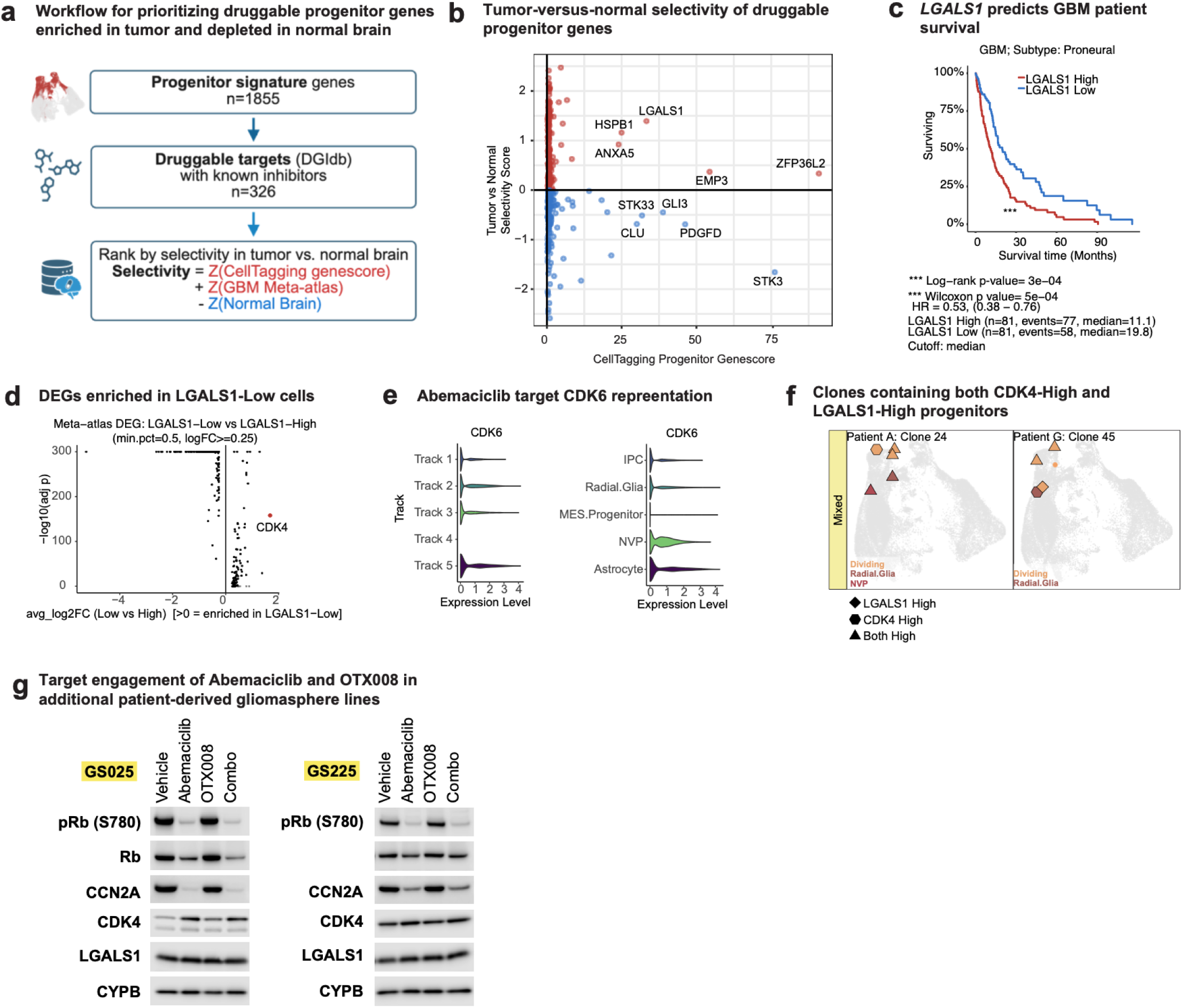
Identification of progenitor-associated drug targets with tumor-selective expression a) Strategy used to identify candidate therapeutic targets from lineage-informed progenitor programs. Progenitor signature genes (n = 1,855) were first defined from the CellTagging dataset based on differential expression between progenitor and non-progenitor malignant cells. These genes were queried against the Drug-Gene Interaction Database (DGIdb) to identify targets with known inhibitor interactions, yielding 326 druggable genes. Druggable candidates were then ranked by tumor selectivity using a composite score that integrates progenitor specificity from the CellTagging genescore together with relative expression in a primary GBM meta-atlas and depletion in a normal adult brain meta-atlas, enabling prioritization of progenitor targets with maximal therapeutic window. b) Scatter plot shows progenitor-associated druggable genes identified by querying the Drug-Gene Interaction Database (DGIdb) for inhibitor-linked interactions among progenitor signature genes. The x-axis shows the CellTagging progenitor genescore, a composite metric reflecting both specificity and enrichment in progenitor cells, calculated as the ratio of the fraction of progenitor cells expressing a gene to the fraction of non-progenitor cells expressing the gene, multiplied by the average log_2_ fold change. The y-axis shows a tumor-normal score calculated as the difference between z-scored mean expression in a GBM meta-atlas and z-scored mean expression in a normal adult brain meta-atlas. Each point represents a gene. Genes with positive tumor-normal scores (red) are enriched in tumor relative to normal brain, whereas genes with negative scores (blue) show higher relative expression in normal tissue. c) Kaplan-Meier survival curves for proneural GBM patients in the TCGA dataset stratified by LGALS1 expression using a median cutoff. Patients with LGALS1 High tumors (red; n = 81, events = 77, median survival = 11.1 months) exhibit reduced overall survival compared to the LGALS1 Low group (blue; n = 81, events = 58, median survival = 19.8 months). Significance was assessed by log-rank test (p = 3e−04) and Wilcoxon test (p = 5e−04); hazard ratio (HR) = 0.53 (95% CI, 0.38-0.76). d) Volcano plot summarizing differential expression in the primary GBM meta-atlas comparing LGALS1-Low versus LGALS1-High cells. e) Differential expression in the primary GBM meta-atlas comparing LGALS1-Low vs LGALS1-High cells (Low vs High). Each point is a gene (x-axis, avg_log2FC Low vs High; y-axis, −log10 adjusted P value). Positive fold-change indicates genes enriched in LGALS1-Low cells; CDK4 is highlighted/labeled. Differential expression used a Wilcoxon test with min.pct = 0.5 and logFC threshold = 0.25. f) Representative immunoblots from two additional patient-derived gliomasphere lines (GS025 and GS225) treated for 7 days with Vehicle (DMSO), Abemaciclib (1 µM), OTX008 (2.5 µM), or the combination.

**Supplementary Figure 6.**
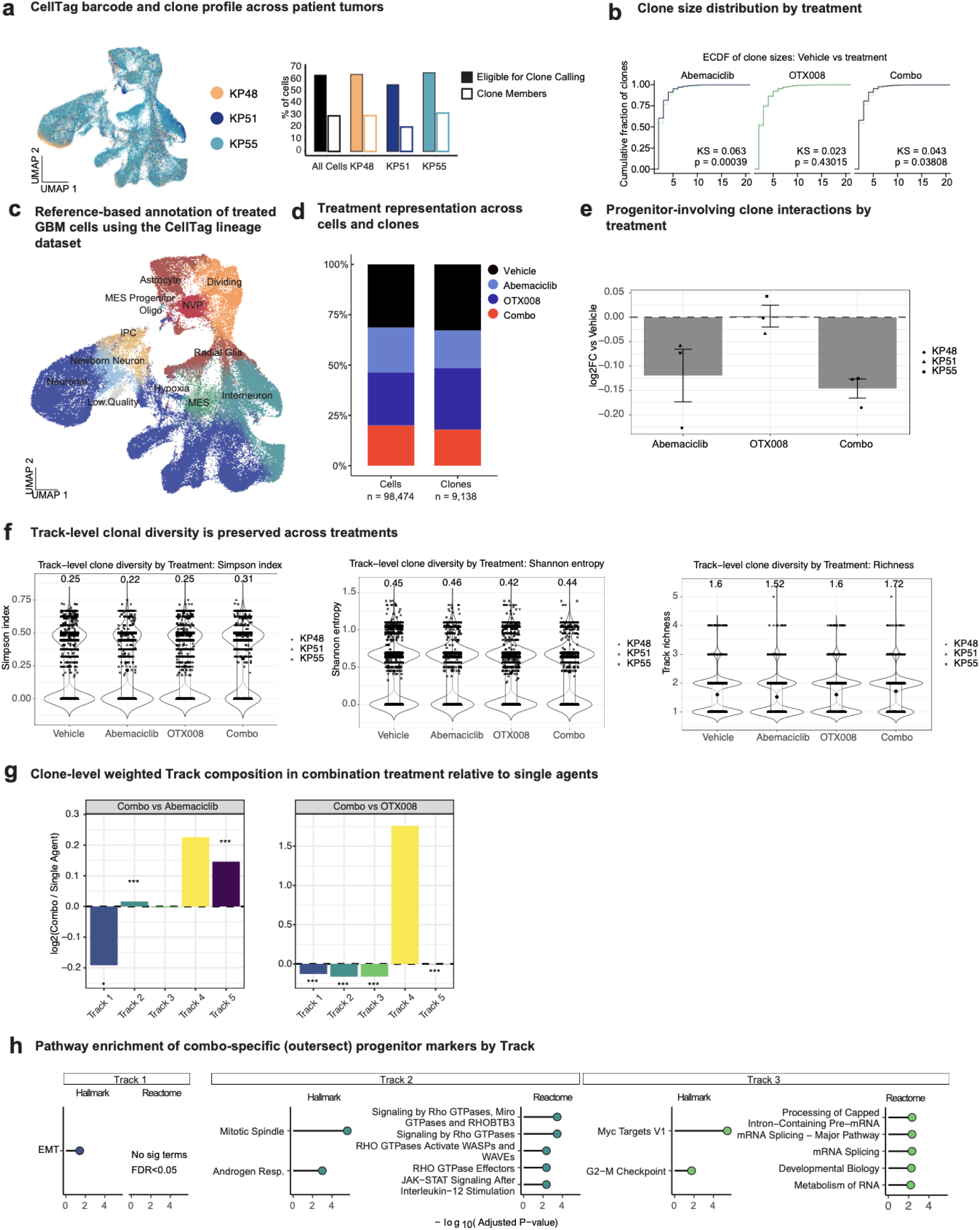
Quality control and extended analyses supporting treatment effects on progenitor tracks a) UMAP visualization of CellTag-labeled GBM cells from three primary IDH1^WT^ GBM tumors (KP48, KP51, KP55), colored by patient of origin. Right, bar plots showing the fraction of all cells eligible for clone calling based on CellTag barcode complexity (filled bars) and the fraction assigned as clone members (outlined bars) for each tumor. Percentages are calculated using all quality-controlled malignant cells as the denominator, illustrating comparable lineage-tracing coverage across patients. b) Empirical cumulative distribution functions (ECDFs) of inferred clone sizes (cells per clone) comparing Vehicle to each treatment condition (Abemaciclib, OTX008, Combo). For each facet, clone sizes were computed as the number of cells assigned to each Clone_ID within that treatment, and Vehicle clone sizes were overlaid for direct comparison. Kolmogorov-Smirnov (KS) statistics and two-sided p-values are shown for Vehicle versus each treatment. c) UMAP of GBM cells following reference-based mapping to the CellTag lineage-tracing dataset shown in Fig. 1c. Cell identities were assigned using Seurat MapQuery, enabling consistent annotation of progenitor and differentiated cell states across treatment conditions. Annotated populations include progenitor states (Radial Glia, IPC, MES.Progenitor, Astrocyte, NVP) and differentiated or state-defined populations (Neuronal, Newborn Neuron, Interneuron, Oligodendrocyte-like, MES, Hypoxic, Dividing). d) Fraction of all cells (left) and inferred clones (right) assigned to each treatment condition. e) Vehicle-normalized change in the proportion of clone interaction pairs involving at least one progenitor cell. Clone interactions were defined as unordered cell type co-occurrence pairs present within each clone (binary per clone), computed using clone-member cells and restricted to clones containing ≥1 progenitor cell. Bars show the mean per-patient log₂ fold change versus Vehicle (log₂FC), with error bars indicating ± s.e.m. across patients (KP48, KP51, KP55); points denote individual patient values. f) Track-level clone diversity across treatment conditions, quantified using three complementary metrics: Simpson index (left), Shannon entropy (middle), and Track richness (right). Diversity metrics were computed per inferred clone using clone-member cells, with Tracks assigned by mapping each cell type to its Track category (Methods). Violin plots show the distribution of per-clone values, with overlaid boxplots indicating median and interquartile range; points denote individual clones (shapes indicate patient of origin). Statistical testing was performed using patient-level replication: for each patient and treatment, clone-level metrics were summarized as the median across that patient’s clones and compared to Vehicle using paired Wilcoxon signed-rank tests with Benjamini-Hochberg correction across all metric × treatment comparisons. All comparisons were non-significant (FDR = 1), indicating no detectable treatment-associated change in track-level clonal diversity. g) Weighted log₂ fold change in clone-member Track representation comparing Combo vs Abemaciclib (left) and Combo vs OTX008 (right). For each treatment arm, clone-member Track proportions were weighted by the inverse of baseline Track frequency within that treatment (1 / p(Track | Treatment)) to normalize for global Track abundance differences. Log₂ fold change was computed as log₂(Combo / Single Agent) using weighted proportions. Statistical significance was assessed using clone-level replication with two-sided one-sample Wilcoxon tests against 0, followed by Benjamini–Hochberg correction across all Track × contrast comparisons (* FDR < 0.05, ** FDR < 0.01, *** FDR < 0.001). Positive values indicate relative enrichment in Combo-treated clones, whereas negative values indicate depletion relative to the corresponding single agent. h) Gene set enrichment of Track-stratified progenitor “outersect” markers. Progenitor cells were grouped by Track and differential expression was performed testing Vehicle vs Abemaciclib, OTX008, and Combo. For each Track, the outersect gene set was defined as genes significant in Vehicle vs Combo (p_adj < 0.05) minus the union of genes significant in Vehicle vs Abemaciclib or Vehicle vs OTX008. Lollipop plots show the top five enriched Hallmark and Reactome terms per Track (FDR < 0.05), ranked by adjusted P value (x-axis, −log10 adjusted P value); panels with no significant terms are labeled.

## Bibliography

1. Lan, X., Jorg, D.J., Cavalli, F.M.G., Richards, L.M., Nguyen, L.V., Vanner, R.J., Guilhamon, P., Lee, L., Kushida, M.M., Pellacani, D., et al. (2017). Fate mapping of human glioblastoma reveals an invariant stem cell hierarchy. Nature 549, 227–232. 10.1038/nature23666.

2. Neftel, C., Laffy, J., Filbin, M.G., Hara, T., Shore, M.E., Rahme, G.J., Richman, A.R., Silverbush, D., Shaw, M.L., Hebert, C.M., et al. (2019). An Integrative Model of Cellular States, Plasticity, and Genetics for Glioblastoma. Cell 178, 835–849.e21. 10.1016/j.cell.2019.06.024.

3. Alcantara Llaguno, S.R., Wang, Z., Sun, D., Chen, J., Xu, J., Kim, E., Hatanpaa, K.J., Raisanen, J.M., Burns, D.K., Johnson, J.E., et al. (2015). Adult Lineage Restricted CNS Progenitors Specify Distinct Glioblastoma Subtypes. Cancer Cell 28, 429–440. 10.1016/j.ccell.2015.09.007.

4. Wang, Z., Sun, D., Chen, Y.-J., Xie, X., Shi, Y., Tabar, V., Brennan, C.W., Bale, T.A., Jayewickreme, C.D., Laks, D.R., et al. (2020). Cell lineage-based stratification for glioblastoma. Cancer Cell 38, 366–379.e8. 10.1016/j.ccell.2020.06.003.

5. Liau, B.B., Sievers, C., Donohue, L.K., Gillespie, S.M., Flavahan, W.A., Miller, T.E., Venteicher, A.S., Hebert, C.H., Carey, C.D., Rodig, S.J., et al. (2017). Adaptive Chromatin Remodeling Drives Glioblastoma Stem Cell Plasticity and Drug Tolerance. Cell Stem Cell 20, 233–246.e7. 10.1016/j.stem.2016.11.003.

6. Saraswat, M., Rueda-Gensini, L., Heinzelmann, E., Gracia, T., Memi, F., de Jong, G., Straub, J., Schloo, C., Hoffmann, D.C., Jung, E., et al. (2025). Decoding Plasticity Regulators and Transition Trajectories in Glioblastoma with Single-cell Multiomics. Preprint at bioRxiv, 10.1101/2025.05.13.653733.

7. Eyler, C.E., Matsunaga, H., Hovestadt, V., Vantine, S.J., van Galen, P., and Bernstein, B.E. (2020). Single-cell lineage analysis reveals genetic and epigenetic interplay in glioblastoma drug resistance. Genome Biol 21, 174. 10.1186/s13059-020-02085-1.

8. Wang, R., Sharma, R., Shen, X., Laughney, A.M., Funato, K., Clark, P.J., Shpokayte, M., Morgenstern, P., Navare, M., Xu, Y., et al. (2020). Adult Human Glioblastomas Harbor Radial Glia-like Cells. Stem Cell Reports 14, 338–350. 10.1016/j.stemcr.2020.01.007.

9. Bhaduri, A., Di Lullo, E., Jung, D., Müller, S., Crouch, E.E., Espinosa, C.S., Ozawa, T., Alvarado, B., Spatazza, J., Cadwell, C.R., et al. (2020). Outer Radial Glia-like Cancer Stem Cells Contribute to Heterogeneity of Glioblastoma. Cell Stem Cell 26, 48–63.e6. 10.1016/j.stem.2019.11.015.

10. Couturier, C.P., Ayyadhury, S., Le, P.U., Nadaf, J., Monlong, J., Riva, G., Allache, R., Baig, S., Yan, X., Bourgey, M., et al. (2020). Single-cell RNA-seq reveals that glioblastoma recapitulates a normal neurodevelopmental hierarchy. Nat Commun 11, 3406. 10.1038/s41467-020-17186-5.

11. Wang, L., Wang, C., Moriano, J.A., Chen, S., Zuo, G., Cebrian-Silla, A., Zhang, S., Mukhtar, T., Wang, S., Song, M., et al. (2025). Molecular and cellular dynamics of the developing human neocortex. Nature. 10.1038/s41586-024-08351-7.

12. Wang, L., Babikir, H., Muller, S., Yagnik, G., Shamardani, K., Catalan, F., Kohanbash, G., Alvarado, B., Di Lullo, E., Kriegstein, A., et al. (2019). The Phenotypes of Proliferating Glioblastoma Cells Reside on a Single Axis of Variation. Cancer Discov 9, 1708–1719. 10.1158/2159-8290.CD-19-0329.

13. Yu, K., Hu, Y., Wu, F., Guo, Q., Qian, Z., Hu, W., Chen, J., Wang, K., Fan, X., Wu, X., et al. (2020). Surveying brain tumor heterogeneity by single-cell RNA-sequencing of multi-sector biopsies. National Science Review 7, 1306–1318. 10.1093/nsr/nwaa099.

14. Chen, J., Li, Y., Yu, T.-S., McKay, R.M., Burns, D.K., Kernie, S.G., and Parada, L.F. (2012). A restricted cell population propagates glioblastoma growth after chemotherapy. Nature 488, 522–526. 10.1038/nature11287.

15. Pollen, A.A., Nowakowski, T.J., Chen, J., Retallack, H., Sandoval-Espinosa, C., Nicholas, C.R., Shuga, J., Liu, S.J., Oldham, M.C., Diaz, A., et al. (2015). Molecular identity of human outer radial glia during cortical development. Cell 163, 55–67. 10.1016/j.cell.2015.09.004.

16. Darmanis, S., Sloan, S.A., Croote, D., Mignardi, M., Chernikova, S., Samghababi, P., Zhang, Y., Neff, N., Kowarsky, M., Caneda, C., et al. (2017). Single-Cell RNA-Seq Analysis of Infiltrating Neoplastic Cells at the Migrating Front of Human Glioblastoma. Cell Reports 21, 1399–1410. 10.1016/j.celrep.2017.10.030.

17. Mathur, R., Wang, Q., Schupp, P.G., Nikolic, A., Hilz, S., Hong, C., Grishanina, N.R., Kwok, D., Stevers, N.O., Jin, Q., et al. (2024). Glioblastoma evolution and heterogeneity from a 3D whole-tumor perspective. Cell 187, 446–463.e16. 10.1016/j.cell.2023.12.013.

18. Nomura, M., Spitzer, A., Johnson, K.C., Garofano, L., Nehar-Belaid, D., Galili Darnell, N., Greenwald, A.C., Bussema, L., Oh, Y.T., Varn, F.S., et al. (2025). The multilayered transcriptional architecture of glioblastoma ecosystems. Nat Genet 57, 1155–1167. 10.1038/s41588-025-02167-5.

19. Patel, A.P., Tirosh, I., Trombetta, J.J., Shalek, A.K., Gillespie, S.M., Wakimoto, H., Cahill, D.P., Nahed, B.V., Curry, W.T., Martuza, R.L., et al. (2014). Single-cell RNA-seq highlights intratumoral heterogeneity in primary glioblastoma. Science 344, 1396–1401. 10.1126/science.1254257.

20. Ruiz-Moreno, C., Salas, S.M., Samuelsson, E., Minaeva, M., Ibarra, I., Grillo, M., Brandner, S., Roy, A., Forsberg-Nilsson, K., Kranendonk, M.E.G., et al. (2025). Charting the single-cell and spatial landscape of IDH-wild-type glioblastoma with GBmap. Neuro Oncol 27, 2281–2295. 10.1093/neuonc/noaf113.

21. Jacob, F., Salinas, R.D., Zhang, D.Y., Nguyen, P.T.T., Schnoll, J.G., Wong, S.Z.H., Thokala, R., Sheikh, S., Saxena, D., Prokop, S., et al. (2020). A Patient-Derived Glioblastoma Organoid Model and Biobank Recapitulates Inter- and Intra-tumoral Heterogeneity. Cell 180, 188–204.e22. 10.1016/j.cell.2019.11.036.

22. Mangena, V., Chanoch-Myers, R., Sartore, R., Paulsen, B., Gritsch, S., Weisman, H., Hara, T., Breakefield, X.O., Breyne, K., Regev, A., et al. (2025). Glioblastoma Cortical Organoids Recapitulate Cell-State Heterogeneity and Intercellular Transfer. Cancer Discov 15, 299–315. 10.1158/2159-8290.CD-23-1336.

23. Peng, T., Ma, X., Hua, W., Wang, C., Chu, Y., Sun, M., Fermi, V., Hamelmann, S., Lindner, K., Shao, C., et al. (2025). Individualized patient tumor organoids faithfully preserve human brain tumor ecosystems and predict patient response to therapy. Cell Stem Cell 32, 652–669 e11. 10.1016/j.stem.2025.01.002.

24. Shimizu, F., Hovinga, K.E., Metzner, M., Soulet, D., and Tabar, V. (2011). Organotypic explant culture of glioblastoma multiforme and subsequent single-cell suspension. Curr Protoc Stem Cell Biol Chapter 3, Unit3 5. 10.1002/9780470151808.sc0305s19.

25. Ge, W., Kan, R.L., Yilgor, C., Fazzari, E., Nano, P.R., Azizad, D.J., Shinglot, H., Li, M., Ito, J.Y., Tse, C., et al. (2026). Human organoid tumor transplantation identifies functional glioblastoma-microenvironment communication mediated by PTPRZ1. Cell Rep 45, 116848. 10.1016/j.celrep.2025.116848.

26. Biddy, B.A., Kong, W., Kamimoto, K., Guo, C., Waye, S.E., Sun, T., and Morris, S.A. (2018). Single-cell mapping of lineage and identity in direct reprogramming. Nature 564, 219–224. 10.1038/s41586-018-0744-4.

27. Fazzari, E., Azizad, D.J., Yu, K., Ge, W., Li, M.X., Nano, P.R., Kan, R.L., Tum, H.A., Tse, C., Bayley, N.A., et al. (2024). Glioblastoma Neurovascular Progenitor Orchestrates Tumor Cell Type Diversity. bioRxiv. 10.1101/2024.07.24.604840.

28. Sloan, S.A., Darmanis, S., Huber, N., Khan, T.A., Birey, F., Caneda, C., Reimer, R., Quake, S.R., Barres, B.A., and Paşca, S.P. (2017). Human Astrocyte Maturation Captured in 3D Cerebral Cortical Spheroids Derived from Pluripotent Stem Cells. Neuron 95, 779–790.e6. 10.1016/j.neuron.2017.07.035.

29. Sojka, C., Wang, H.-L.V., Bhatia, T.N., Li, Y., Chopra, P., Sing, A., Voss, A., King, A., Wang, F., Joseph, K., et al. (2025). Mapping the developmental trajectory of human astrocytes reveals divergence in glioblastoma. Nat Cell Biol 27, 347–359. 10.1038/s41556-024-01583-9.

30. Hara, T., Chanoch-Myers, R., Mathewson, N.D., Myskiw, C., Atta, L., Bussema, L., Eichhorn, S.W., Greenwald, A.C., Kinker, G.S., Rodman, C., et al. (2021). Interactions between cancer cells and immune cells drive transitions to mesenchymal-like states in glioblastoma. Cancer Cell 39, 779–792.e11. 10.1016/j.ccell.2021.05.002.

31. Ravi, V.M., Will, P., Kueckelhaus, J., Sun, N., Joseph, K., Salie, H., Vollmer, L., Kuliesiute, U., von Ehr, J., Benotmane, J.K., et al. (2022). Spatially resolved multi-omics deciphers bidirectional tumor-host interdependence in glioblastoma. Cancer Cell 40, 639–655 e13. 10.1016/j.ccell.2022.05.009.

32. Greenwald, A.C., Darnell, N.G., Hoefflin, R., Simkin, D., Mount, C.W., Gonzalez Castro, L.N., Harnik, Y., Dumont, S., Hirsch, D., Nomura, M., et al. (2024). Integrative spatial analysis reveals a multi-layered organization of glioblastoma. Cell 187, 2485–2501 e26. 10.1016/j.cell.2024.03.029.

33. Delgado, R.N., Allen, D.E., Keefe, M.G., Mancia Leon, W.R., Ziffra, R.S., Crouch, E.E., Alvarez-Buylla, A., and Nowakowski, T.J. (2022). Individual human cortical progenitors can produce excitatory and inhibitory neurons. Nature 601, 397–403. 10.1038/s41586-021-04230-7.

34. Cannon, M., Stevenson, J., Stahl, K., Basu, R., Coffman, A., Kiwala, S., McMichael, J.F., Kuzma, K., Morrissey, D., Cotto, K., et al. (2024). DGIdb 5.0: rebuilding the drug–gene interaction database for precision medicine and drug discovery platforms. Nucleic Acids Res 52, D1227–D1235. 10.1093/nar/gkad1040.

35. Guda, M.R., Tsung, A.J., Asuthkar, S., and Velpula, K.K. (2022). Galectin-1 activates carbonic anhydrase IX and modulates glioma metabolism. Cell Death Dis 13, 574. 10.1038/s41419-022-05024-z.

36. Imaizumi, Y., Sakaguchi, M., Morishita, T., Ito, M., Poirier, F., Sawamoto, K., and Okano, H. (2011). Galectin-1 is expressed in early-type neural progenitor cells and down-regulates neurogenesis in the adult hippocampus. Mol Brain 4, 7. 10.1186/1756-6606-4-7.

37. Sakaguchi, M., Shingo, T., Shimazaki, T., Okano, H.J., Shiwa, M., Ishibashi, S., Oguro, H., Ninomiya, M., Kadoya, T., Horie, H., et al. (2006). A carbohydrate-binding protein, Galectin-1, promotes proliferation of adult neural stem cells. Proc Natl Acad Sci U S A 103, 7112–7117. 10.1073/pnas.0508793103.

38. Sharanek, A., Burban, A., Hernandez-Corchado, A., Madrigal, A., Fatakdawala, I., Najafabadi, H.S., Soleimani, V.D., and Jahani-Asl, A. (2021). Transcriptional control of brain tumor stem cells by a carbohydrate binding protein. Cell Rep 36, 109647. 10.1016/j.celrep.2021.109647.

39. Rahman, R., Trippa, L., Lee, E.Q., Arrillaga-Romany, I., Fell, G., Touat, M., McCluskey, C., Wiley, J., Gaffey, S., Drappatz, J., et al. (2023). Inaugural Results of the Individualized Screening Trial of Innovative Glioblastoma Therapy: A Phase II Platform Trial for Newly Diagnosed Glioblastoma Using Bayesian Adaptive Randomization. J Clin Oncol 41, 5524–5535. 10.1200/JCO.23.00493.

40. Taylor, J.W., Parikh, M., Phillips, J.J., James, C.D., Molinaro, A.M., Butowski, N.A., Clarke, J.L., Oberheim-Bush, N.A., Chang, S.M., Berger, M.S., et al. (2018). Phase-2 trial of palbociclib in adult patients with recurrent RB1-positive glioblastoma. J Neurooncol 140, 477–483. 10.1007/s11060-018-2977-3.

41. Tannous, B.A. (2009). Gaussia luciferase reporter assay for monitoring biological processes in culture and in vivo. Nat Protoc 4, 582–591. 10.1038/nprot.2009.28.

42. Dirkse, A., Golebiewska, A., Buder, T., Nazarov, P.V., Muller, A., Poovathingal, S., Brons, N.H.C., Leite, S., Sauvageot, N., Sarkisjan, D., et al. (2019). Stem cell-associated heterogeneity in Glioblastoma results from intrinsic tumor plasticity shaped by the microenvironment. Nat Commun 10, 1787. 10.1038/s41467-019-09853-z.

43. Larsson, I., Dalmo, E., Elgendy, R., Niklasson, M., Doroszko, M., Segerman, A., Jörnsten, R., Westermark, B., and Nelander, S. (2021). Modeling glioblastoma heterogeneity as a dynamic network of cell states. Mol Syst Biol 17, MSB202010105. 10.15252/msb.202010105.

44. Singh, S.K., Hawkins, C., Clarke, I.D., Squire, J.A., Bayani, J., Hide, T., Henkelman, R.M., Cusimano, M.D., and Dirks, P.B. (2004). Identification of human brain tumour initiating cells. Nature 432, 396–401. 10.1038/nature03128.

45. Lee, E.Q., Trippa, L., Fell, G., Rahman, R., Arrillaga-Romany, I., Touat, M., Drappatz, J., Welch, M.R., Galanis, E., Ahluwalia, M.S., et al. (2021). Preliminary results of the abemaciclib arm in the Individualized Screening Trial of Innovative Glioblastoma Therapy (INSIGhT): A phase II platform trial using Bayesian adaptive randomization. J Clin Oncol 39, 2014–2014. 10.1200/JCO.2021.39.15_suppl.2014.

46. Bhaduri, A., Andrews, M.G., Mancia Leon, W., Jung, D., Shin, D., Allen, D., Jung, D., Schmunk, G., Haeussler, M., Salma, J., et al. (2020). Cell stress in cortical organoids impairs molecular subtype specification. Nature 578, 142–148. 10.1038/s41586-020-1962-0.

47. Pollen, A.A., Bhaduri, A., Andrews, M.G., Nowakowski, T.J., Meyerson, O.S., Mostajo-Radji, M.A., Di Lullo, E., Alvarado, B., Bedolli, M., Dougherty, M.L., et al. (2019). Establishing Cerebral Organoids as Models of Human-Specific Brain Evolution. Cell 176, 743–756 e17. 10.1016/j.cell.2019.01.017.

48. Velasco, S., Kedaigle, A.J., Simmons, S.K., Nash, A., Rocha, M., Quadrato, G., Paulsen, B., Nguyen, L., Adiconis, X., Regev, A., et al. (2019). Individual brain organoids reproducibly form cell diversity of the human cerebral cortex. Nature 570, 523–527. 10.1038/s41586-019-1289-x.

49. Hao, Y., Stuart, T., Kowalski, M.H., Choudhary, S., Hoffman, P., Hartman, A., Srivastava, A., Molla, G., Madad, S., Fernandez-Granda, C., et al. (2024). Dictionary learning for integrative, multimodal and scalable single-cell analysis. Nat Biotechnol 42, 293–304. 10.1038/s41587-023-01767-y.

50. Shekhar, K., Lapan, S.W., Whitney, I.E., Tran, N.M., Macosko, E.Z., Kowalczyk, M., Adiconis, X., Levin, J.Z., Nemesh, J., Goldman, M., et al. (2016). Comprehensive Classification of Retinal Bipolar Neurons by Single-Cell Transcriptomics. Cell 166, 1308–1323.e30. 10.1016/j.cell.2016.07.054.

51. McGinnis, C.S., Murrow, L.M., and Gartner, Z.J. (2019). DoubletFinder: Doublet Detection in Single-Cell RNA Sequencing Data Using Artificial Nearest Neighbors. Cell Syst 8, 329–337 e4. 10.1016/j.cels.2019.03.003.

52. Home · broadinstitute/infercnv Wiki https://github.com/broadinstitute/inferCNV/wiki.

53. Nano, P.R., Fazzari, E., Azizad, D., Martija, A., Nguyen, C.V., Wang, S., Giang, V., Kan, R.L., Yoo, J., Wick, B., et al. (2025). Integrated analysis of molecular atlases unveils modules driving developmental cell subtype specification in the human cortex. Nat Neurosci 28, 949–963. 10.1038/s41593-025-01933-2.

54. Nowakowski, T.J., Bhaduri, A., Pollen, A.A., Alvarado, B., Mostajo-Radji, M.A., Di Lullo, E., Haeussler, M., Sandoval-Espinosa, C., Liu, S.J., Velmeshev, D., et al. (2017). Spatiotemporal gene expression trajectories reveal developmental hierarchies of the human cortex. Science 358, 1318–1323. 10.1126/science.aap8809.

55. Tsyben, A., Dannhorn, A., Hamm, G., Pitoulias, M., Couturier, D.-L., Sawle, A., Briggs, M., Wright, A.J., Brodie, C., Mendil, L., et al. (2025). Cell-intrinsic metabolic phenotypes identified in patients with glioblastoma, using mass spectrometry imaging of 13C-labelled glucose metabolism. Nat Metab 7, 928–939. 10.1038/s42255-025-01293-y.

